# Aberrant claudin-6–adhesion signal promotes endometrial cancer progression via estrogen receptor α

**DOI:** 10.1101/2020.05.15.097659

**Authors:** Manabu Kojima, Kotaro Sugimoto, Mizuko Tanaka, Yuta Endo, Naoki Ichikawa-Tomikawa, Korehito Kashiwagi, Hitomi Kato, Tsuyoshi Honda, Shigenori Furukawa, Hiroshi Nishiyama, Takafumi Watanabe, Shu Soeda, Keiya Fujimori, Hideki Chiba

## Abstract

Cell adhesion proteins not only maintain tissue integrity but also possess signaling abilities to organize diverse cellular events in physiological and pathological processes; however, the underlying mechanism remains obscure. Among cell adhesion molecules, the claudin (CLDN) family often possesses aberrant expression in various cancers, but the biological relevance and molecular basis have not yet been established. Here, we show that high CLDN6 expression promotes endometrial cancer progression and represents the poor prognostic marker. The second extracellular domain and Y196/200 of CLDN6 were required to recruit and activate Src-family kinases (SFKs) and to stimulate malignant phenotypes. Importantly, we demonstrate that the CLDN6/SFK/PI3K-dependent AKT and SGK (serum- and glucocorticoid-regulated kinase) signalings target Ser518 in the human estrogen receptor α and ligand-independently activate target genes in endometrial cancer cells, resulting in cancer development. The identification of this machinery highlights regulation of the transcription factors by cell adhesion to advance tumor progression.

## Introduction

Endometrial cancer represents the most common gynecological malignancy in developed countries, with an increased prevalence worldwide (1). Although it has been considered to occur during the postmenopausal period, cases diagnosed in premenopausal women are growing (2). The risk factors for endometrial cancer include an excess of endogenous and exogenous estrogens, older age, obesity, and nulliparity (3,4). Patients with endometrial cancer are often found at the early stages and possess a relatively favorable prognosis. However, up to 20% of cases recur after primary surgery, and the 5-year overall survival rates for the International Federation of Gynecology and Obstetrics (FIGO) stages III and IV are 57–66% and 20–26%, respectively (5). Therefore, biomarkers that reflect the malignant behavior of endometrial cancer are required to identify patients with poor outcome.

Claudins (CLDNs) are major proteins of tight junctions, the apical-most components of apical junctional complexes (6–9). The CLDN family is composed of 24 members in humans, and displays distinct expression patterns in tissue- and cell-type selective manners. CLDNs also show aberrant expression in a variety of cancer tissues (10). These tetraspanning membrane proteins have a short cytoplasmic N-terminus, two extracellular loops (EC1 and EC2) and a C-terminal cytoplasmic domain. CLDNs act as paracellular barriers or pores via the EC1 to regulate selective transport of ions and substances. On the other hand, CLDN-EC2 participates not only in the binding for *Clostridium Perfringens* enterotoxin (CPE), but also in *trans*-interaction between the plasma membranes of neighboring cells. Furthermore, the C-terminal cytoplasmic domain of CLDNs is thought to propagate intracellular signals, but the underlying molecular basis has not been determined (11).

Among the CLDN family, CLDN6 is expressed in several types of embryonic epithelial cells but not largely in normal adult cells (12–16). In addition, CLDN6 is highly expressed in germ cell tumors, including seminomas, embryonal carcinomas and yolk sac tumors, as well as in some cases of gastric adenocarcinomas, lung adenocarcinomas, ovarian adenocarcinomas and endometrial carcinomas (17,18). However, the biological significance of CLDN6 expression in these cancers remains unclear.

We have recently uncovered that the EC2-dependent engagement of CLDN6 recruits and activates Src-family kinases (SFKs), which in turn phosphorylate CLDN6 at Y196/200 and propagate the PI3K/AKT pathway, and this signaling axis stimulates the retinoic acid receptor γ (RARγ) and estrogen receptor α (ERα) activity (19). Taken together with the notion that ERα acts as a master transcription factor in endometrial cancers (20), we postulated that the CLDN6 signaling modulates the malignant behavior of endometrial cancer cells via ERα. Here, we show that high CLDN6 expression, which expects poor prognosis in endometrial cancer, advances tumor progression. We also demonstrate that the CLDN6/SFK/PI3K axis propagates AKT and SGK (serum- and glucocorticoid-regulated kinase) and targets ERαS518, leading to stimulation of the ERα activity and malignant behaviors in endometrial cancer cells.

## Results

### Establishment of an anti-human CLDN6 mAb

We first generated a novel monoclonal antibody (mAb) against the C-terminal cytoplasmic region of human CLDN6 (*Supplementary Figure S1A*) using the iliac lymph node method (21). Among 384 hybridomas, 24 clones were selected by enzyme-linked immunosorbent assay (ELISA), 20 of which were able to detect CLDN6 by Western blot in HEK293T cells transfected with the corresponding expression vector (*Supplementary Figure S1B and C*). To check the specificity of an anti-human CLDN6 mAb (clone #15) and the previously established anti-mouse CLDN6 polyclonal antibody (pAb; ref. 22), HEK293T cells were transiently transfected with individual CLDN expression vectors, followed by Western blot and immunohistochemical analyses. Clone #15 selectively recognized CLDN6 but not CLDN1, CLDN4, CLDN5 or CLDN9, which are closely related to CLDN6 within the CLDN family (*Supplementary Figure S1D and E*). On the other hand, the anti-CLDN6 pAb reacted not only with CLDN6 but also with overexpressed CLDN4 and CLDN5 to a lesser extent. We also clarified the complementarity-determining regions of clone #15 (*Supplementary Figure S1F*).

### High expression of CLDN6 correlates with poor prognosis in endometrial cancer

Using immunohistochemistry, we next evaluated the expression of CLDN6 in endometrial cancer tissues that resected from 173 patients. Based on semi-quantification using the immunoreactive score, 10 of the 173 cases (5.8%) showed high CLDN6 expression (score 3+). Among the low expression group, 19 (11.0%), 18 (10.4%) and 126 (72.8%) cases had scores 2+, 1+ and 0, respectively. CLDN6 was distributed along the cell membranes of endometrial carcinoma cells (*Figure 1A*). Interestingly, CLDN6 exhibited intratumor heterogeneity, and CLDN6-positive and negative subpopulations were observed in endometrial cancer tissues even in the high CLDN6 expression subjects (*Figure 1B*).

**Figure 1.**
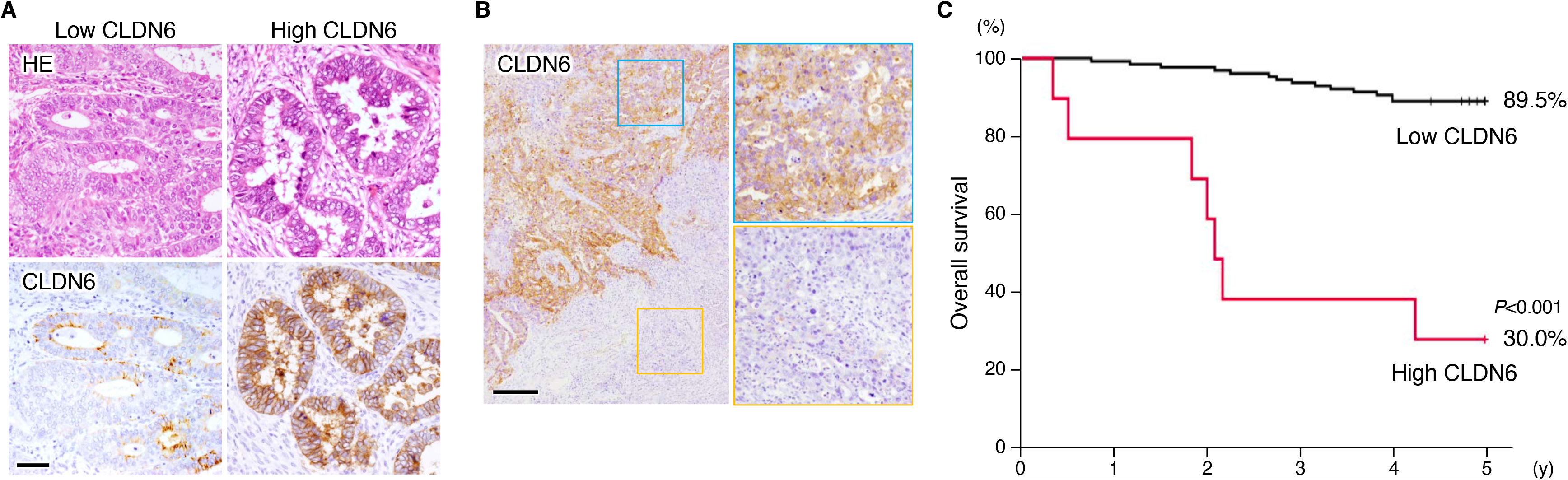
Overexpression of CLDN6 is associated with poor outcome in endometrial cancer patients. (*A*) Representative immunohistological images of high and low CLDN6 expression in endometrial cancer tissues. HE, hematoxylin-eosin. Scale bar, 50 μm. (*B*) Intratumor heterogeneity of CLDN6 protein in the high CLDN6 expression subjects of endometrial cancer. The blue and yellow squares indicate CLDN6-positive and negative subpopulations, respectively. Scale bar, 200 μm. (*C*) Kaplan-Meier plots for high and low CLDN6 expression groups in endometrial cancer subjects.

Kaplan-Meier plots revealed significant differences in overall survival and recurrence-free survival between the two groups (*Figure 1C* and *Supplementary Figure S2*). The five-year survival rate in the high CLDN6 expression group remained at approximately 30%, whereas that in the low expression group was 90%. Among the clinicopathological factors, the high CLDN6 expression was significantly associated with surgical stages III/IV (*P*=0.001), histological grade 3 (*P*=0.004), lymphovascular space involvement (LVSI; *P*=0.001), lymph node metastasis (*P*=0.012) and distant metastasis (*P*=0.014), but not with younger age (*P*=0.122) or histological type (*P*=0.087; *Table 1*). In addition, using the Cox multivariable analysis, stages III/IV (hazard ratio [HR] 10.93, *P*=0.002), distant metastasis (HR 4.68, *P*=0.006) and high CLDN6 expression (HR 3.50, *P*=0.014) possessed independent prognostic variables for overall survival of endometrial cancer patients (*Table 2*).

**Table 1.** Relevance between CLDN6-expression and clinicopathological factors

| parameter | total (n=173) | CLDN6-low<br>(n=163) | CLDN6-high<br>(n=10) | P-value |
| --- | --- | --- | --- | --- |
| Age <50 | 32 (18%) | 32 (20%) | 0 (0%) | 0.122 |
| ≥50 | 141 (32%) | 131 (80%) | 10 (100%) |  |
| Stage I-II | 139 (80%) | 136 (83%) | 3 (30%) | 0.001 |
| III-IV | 34 (20%) | 27 (17%) | 7 (70%) |  |
| Endometrioid | 164 (95%) | 156 (90%) | 8 (80%) | 0.087 |
| Non-endometrioid | 9 (5%) | 7 (10%) | 2 (20%) |  |
| Histological Grade 1-2 | 140 (85%) | 136 (87%) | 4 (50%) | 0.004 |
| 3 | 24 (15%) | 20 (13%) | 4 (50%) |  |
| LVSI (–) | 120 (69%) | 117 (72%) | 3 (30%) | 0.001 |
| LVSI (+) | 53 (31%) | 46 (28%) | 7 (70%) |  |
| N0 | 145 (85%) | 140 (88%) | 5 (50%) | 0.012 |
| N1 | 25 (15%) | 20 (12%) | 5 (50%) |  |
| M0 | 163 (94%) | 156 (96%) | 7 (70%) | 0.014 |
| M1 | 10 (6%) | 7 (4%) | 3 (30%) |  |

LVSI, lymphovascular space involvement; N0/1, negative/positive for lymphnode metastasis; M0/1, negative/positive for distant metastasis.

**Table 2.** Cox multivariable analysis

| variable | HR | 95%CI | P-value |
| --- | --- | --- | --- |
| Age ≥50 | 1.61 | 0.39 – 6.66 | 0.513 |
| Stage III or IV | 10.93 | 2.48 – 48.04 | 0.002 |
| Histological Grade 3 | 2.18 | 0.24 – 3.60 | 0.091 |
| LVSI (+) | 1.91 | 0.51 – 7.18 | 0.340 |
| N1 | 0.45 | 0.13 – 1.61 | 0.220 |
| M1 | 4.68 | 1.57 – 14.01 | 0.006 |
| CLDN6-high | 3.50 | 2.42 – 9.43 | 0.014 |

HR, hazard ratio; CI, confidence interval; LVSI, lymphovascular space involvement; N1, positive for lymphnode metastasis; M1, positive for distant metastasis.

### CLDN6 promotes malignant phenotypes of endometrial carcinoma cells *in vitro* and *in vivo*

We subsequently generated, using the lentiviral vector system, the human endometrial carcinoma cell line Ishikawa expressing CLDN6 (Ishikawa:*CLDN6*; *Figure 2A*). CLDN6 was detected along the cell borders in Ishikawa:*CLDN6* cells, indicating that CLDN6 acted as a cell adhesion molecule (*Figure 2B*). BrdU assay revealed that cellular proliferation was significantly increased in Ishikawa:*CLDN6* cells compared with parental Ishikawa cells (*Figure 2C and D*). In contrast, on the TUNEL assay, few apoptotic cells were observed in both cell lines (*Supplementary Figure S3*). Moreover, wound healing assay demonstrated that cell migration in Ishikawa:*CLDN6* cells was significantly accelerated compared with that in Ishikawa cells (*Figure 2E and F*).

**Figure 2.**
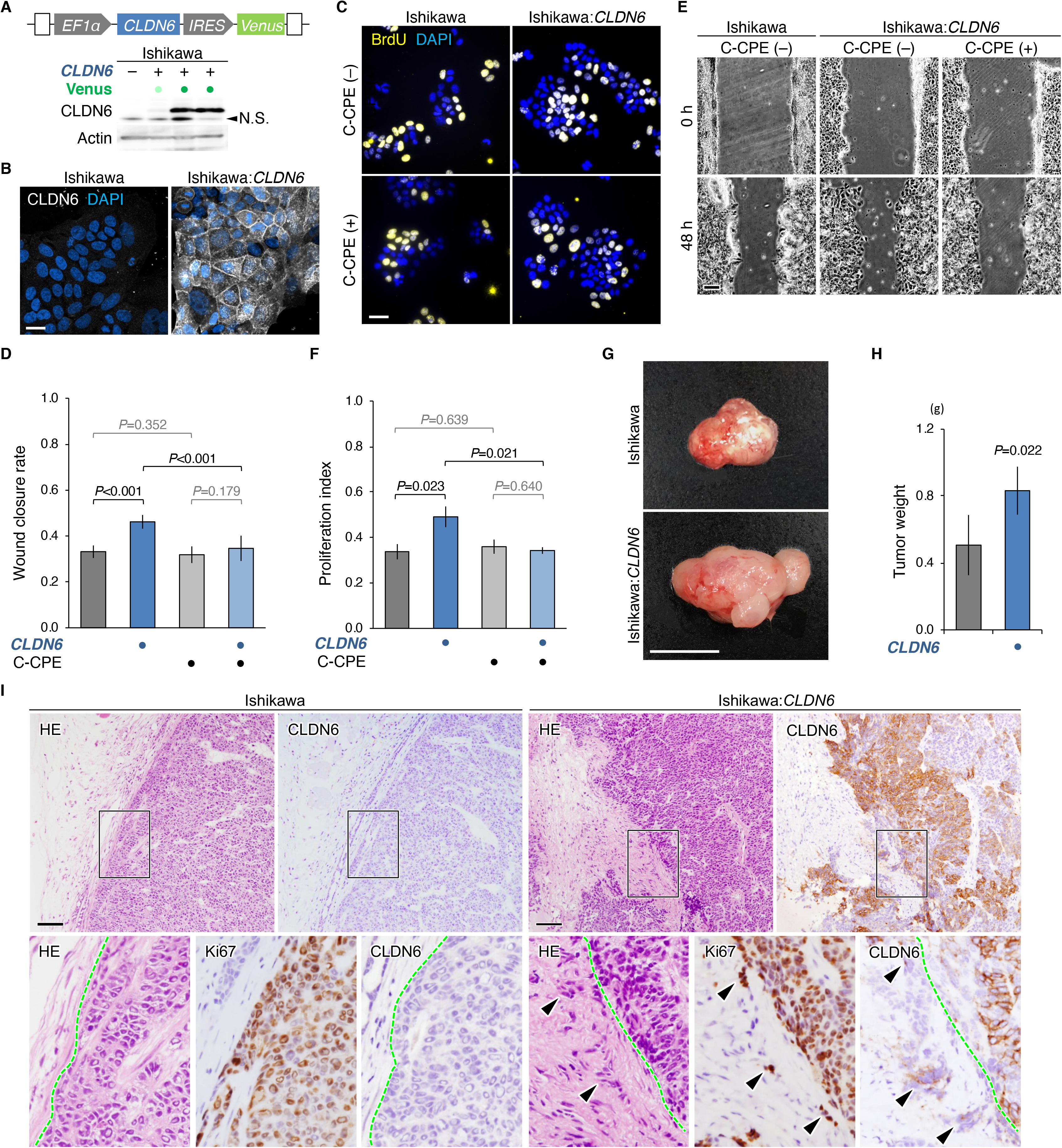
CLDN6 enhances malignant behavior of endometrial carcinoma cells *in vitro* and *in vivo*. (*A* and *B*) Western blot (*A*) and confocal images (*B*) for the indicated proteins in Ishikawa and Ishikawa:*CLDN6* cells. N.S., nonspecific signals. (*C* and *D*) Representative (*C*) and quantitative (*D*) BrdU assay for the indicated cells grown in the presence or absence of 1.0 μg/ml C-CPE. The BrdU/DAPI levels are shown in histograms (mean ± SD; *n* = 6). (*E* and *F*) Typical (*E*) and quantitative (*F*) wound healing assay for the indicated cells grown in the presence or absence of 1.0 μg/ml C-CPE. The values represent wound closure rates (mean ± SD; *n* = 12). Similar results were obtained from another set of experiments for *D* and *F*. (*G*-*I*) Gross and microscopic appearances (*G* and *I*) and weight (*H*) of the indicated xenografts at 28 d after the inoculation. The tumour weight is shown in histograms (mean ± SD; *n* = 4). The regions corresponding the squares include the fibrous capsule around the xenograft tumors, and are enlarged. The boundaries between cancer tissues and the fibrous capsule around the tumor are shown in dashed green lines. Arrowheads indicate invasion into the fibrous capsule around the tumor. Scale bars, 20 μm (*B* and *C*); 50 μm (*E*); 1 cm (*G*); 200 μm (*I*).

We then validated whether the high CLDN6 expression also promoted malignant phenotypes of human endometrial carcinoma cells *in vivo*. Four weeks after inoculation in SCID mice, the tumor weight of Ishikawa:*CLDN6* xenografts was significantly increased compared with that of Ishikawa (*Figure 2G and H*). Neither lymph node nor distant metastasis was grossly evident in these xenografts. Microscopically, Ishikawa:*CLDN6* xenografts were equivalent to Grade 3 endometrial carcinomas that were rich in solid components (*Figure 2I*). Furthermore, intratumor heterogeneity of CLDN6 expression was observed in Ishikawa:*CLDN6* xenograft tissues as in the high CLDN6 expression cases of endometrial cancer subjects. It is also noteworthy that invasion into the fibrous capsule around the tumor was prominent in Ishikawa:*CLDN6* xenografts but hardly in Ishikawa ones.

### The EC2 and Y196/200 of CLDN6 are required for the signaling to activate SFKs in endometrial carcinoma cells and to promote their progression

We next verified the involvement of CLDN6-EC2 and CLDN6-Y196/200 in activation of SFKs and formation of the CLDN6/pSFK complex in human endometrial carcinoma cells. Double immunofluorescence staining showed that pSFK appeared to be concentrated to cell boundaries together with CLDN6 in Ishikawa:*CLDN6* cells (*Figure 3A*). By contrast, pSFK signal was scarcely detected on cell borders in parental Ishikawa cells. When Ishikawa:*CLDN6* cells were exposed to C-terminal half of CPE (C-CPE), which binds to the EC2 of CLDN6 and excludes CLDN6 from cell membranes without alteration in its total protein levels (16,19), the pSFK immunoreactivity was markedly reduced. On Western blot, the levels of pSFK were elevated in Ishikawa:*CLDN6* cells compared with Ishikawa cells, and decreased in both Ishikawa:*CLDN6Y196A* and Ishikawa:*CLDN6Y200A* cells (*Figure 3B*). Immunoprecipitation assay revealed that CLDN6 was associated with pSFK in Ishikawa:*CLDN6* cells, and the CLDN6/pSFK complex was diminished in Ishikawa:*CLDN6* cells on C-CPE treatment as well as in Ishikawa:*CLDN6Y196A* and Ishikawa:*CLDN6Y200A* cells (*Figure 3C and D*).

**Figure 3.**
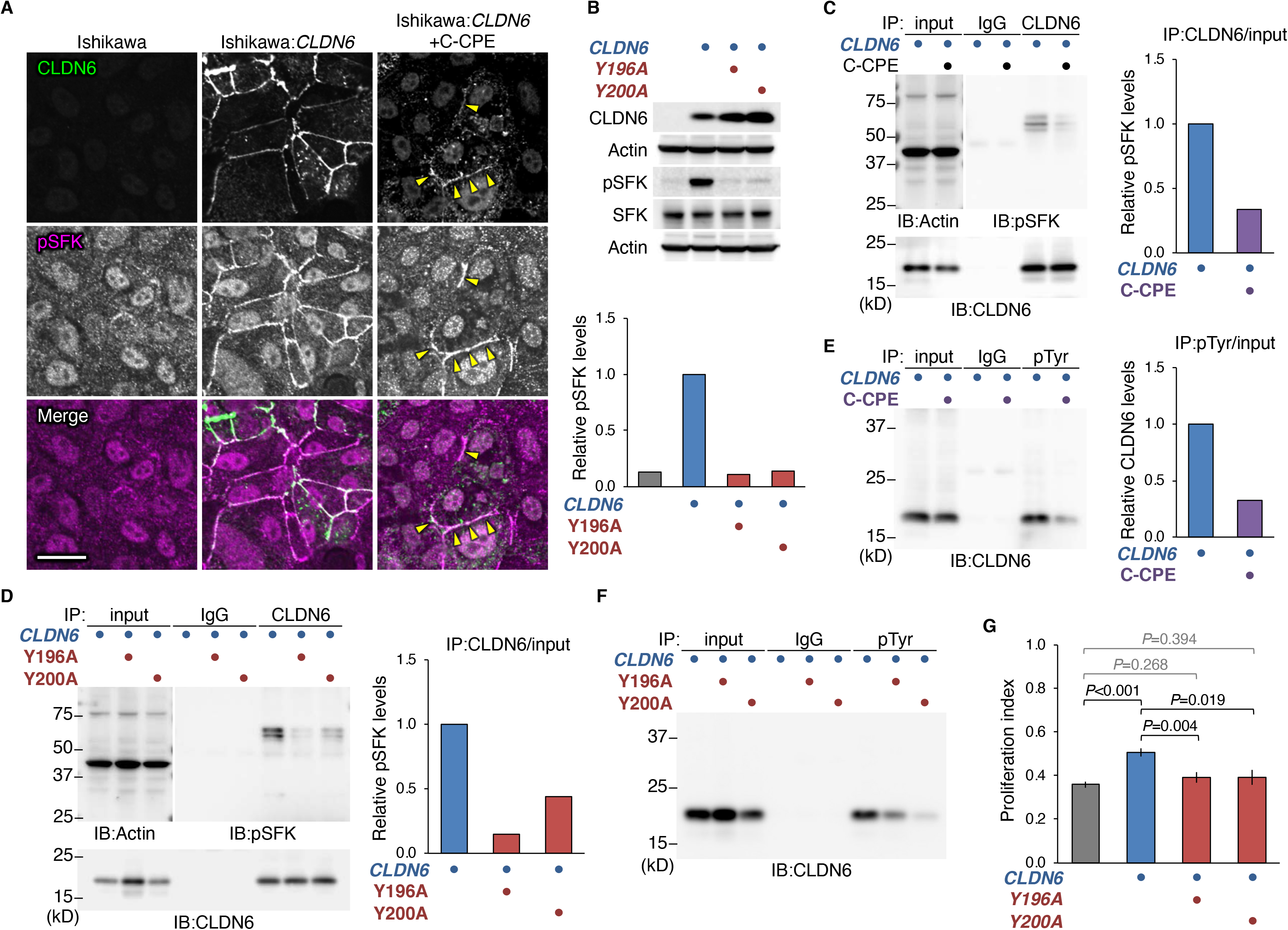
CLDN6 activates SFKs in endometrial carcinoma cells via the EC2 and Y196/200. (*A*) Confocal images for the indicated proteins in Ishikawa and Ishikawa:*CLDN6* cells. Ishikawa:*CLDN6* cells were grown in the presence or absence of 1.0 μg/ml C-CPE. Arrowheads indicate the remaining CLDN6/pSFK signals. Scale bar, 20 μm. (*B*) Western blot for the indicated proteins in the revealed Ishikawa cells. (*C* and *D*) Association between CLDN6 and pSFK in the indicated Ishikawa cell lines. Ishikawa:*CLDN6* cells were exposed to the vehicle or 1.0 μg/ml C-CPE. (*E* and *F*) Tyrosine-phosphorylation of CLDN6 in Ishikawa:*CLDN6* (*E*) and the indicated Ishikawa mutant cells (*F*). Ishikawa:*CLDN6* cells were cultured in the presence or absence of 1.0 μg/ml C-CPE. The protein levels are normalized to the rehybridized actin levels, and the relative levels are shown in the histograms (*B*-*E*). (*G*) BrdU assay for the indicated Ishikawa cells. The BrdU/DAPI levels are shown in histograms (mean ± SD; *n* = 6). Similar results were obtained from another independent experiments.

We also demonstrated that CLDN6 was highly tyrosine-phosphorylated in Ishikawa:*CLDN6* cells, and the phospho-tyrosine levels were suppressed by C-CPE exposure and in both Ishikawa:*CLDN6Y196A* and Ishikawa:*CLDN6Y200A* cells (*Figure 3E and F*). In addition, the promoted cell proliferation and migration in Ishikawa:*CLDN6* cells were significantly reversed by C-CPE treatment (*Figure 2C–F*). Moreover, the CLDN6-enhanced cell proliferation was prevented in Ishikawa:*CLDN6Y196A* or Ishikawa:*CLDN6Y200A* cells (*Figure 3G*). Taken collectively, these results indicated that the CLDN6 signaling activated SFKs and accelerated endometrial cancer progression in the EC2- and Y196/200-dependent manners.

We subsequently validated the involvement of PI3K and the two major downstream cascades AKT and SGK (serum- and glucocorticoid-regulated kinase), which shares the high degree of homology and the same consensus phosphorylation motif (23), in the CLDN6/SFK signaling. To achieve this goal, we used the respective protein kinase inhibitors LY294001, AKT inhibitor VIII and SGK1 inhibitor. As expected, the enhanced cell proliferation in Ishikawa:*CLDN6* cells was prevented by these inhibitors and the SFK inhibitor PP2 (*Supplementary Figure S4*).

### The CLDN6/SFK/PI3K-dependent AKT and SGK signalings target ERα in endometrial carcinoma cells

To evaluate whether the CLDN6-adhesion signaling stimulates the malignant behavior of endometrial carcinoma cells via ERα, we then generated both Ishikawa:*ESR1^−/−^* and Ishikawa:*ESR1^−/−^*:*CLDN6* cells, and compared their phenotypes. We designed the TALEN expression vector that is expected to excise the flanked DNA in exon 2 of *ESR1* genes (*Supplementary Figure S4A*). Knockout of *ESR1* genes was verified by DNA sequence (*Supplementary Figure S4A*), and the lack of ERα protein was confirmed by Western blot and immunostaining (*Supplementary Figure S4B and C*). Note that CLDN6 did not increase cell proliferation or migration capacity in Ishikawa cells in the absence of ERα (*Supplementary Figure S4D–G*).

We also used HEC-1A cells, in which neither CLDN6 nor ERα were expressed, and established cell lines expressing either CLDN6, ERα, or both together (*Figure 4A–C*). Cell growth was significantly elevated in HEC-1A:*ESR1:CLDN6* cells but not in HEC-1A:*CLDN6* or HEC-1A:*ESR1* cells compared with parental HEC-1A cells, (*Figure 4D*). Cell migration was also significantly increased in HEC-1A:*ESR1:CLDN6* cells compared with HEC-1A cells, and was raised in HEC-1A:*CLDN6* and HEC-1A:*ESR1* cells less efficiently than in HEC-1A:*ESR1:CLDN6* cells (*Figure 4E*). Taken together, these results strongly suggested that the CLDN6-adhesion signaling links to ERα in endometrial cancer cells. In addition, exposure of HEC-1A:*ESR1:CLDN6* cells to C-CPE significantly reversed the increase in cell proliferation and migration (*Figure 4D and E*), again indicating the critical role of the EC2 in the CLDN6 signaling.

**Figure 4.**
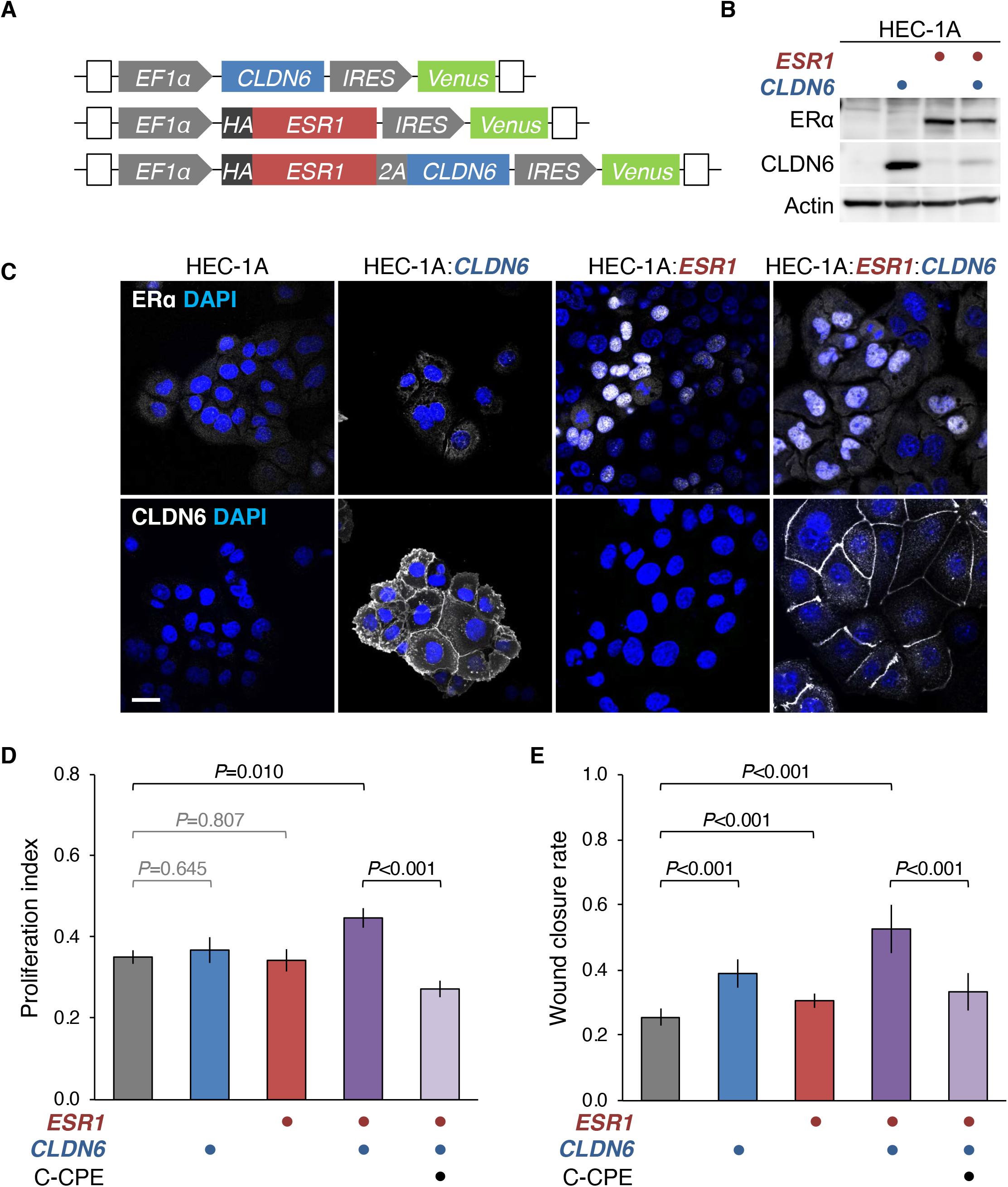
Expression of both CLDN6 and ERα accelerates malignant behavior of HEC-1A cells. (*A*) The construct of *ESR1* and/or *CLDN6* expression vector. EF-1α, elongation factor-1α; IRES, internal ribosome entry site; 2A, self-cleaving peptide. (*B* and *C*) Western blot (*B*) and confocal images (*C*) for the indicated proteins in the revealed cell lines. (*D*) BrdU assay for the indicated cells. The BrdU/DAPI levels are shown in histograms (mean ± SD; *n* = 6). (*E*) Wound healing assay for the indicated cells. The values represent wound closure rates (mean ± SD; *n* = 12). HEC-1A: *ESR1^−/−^*:*CLDN6* cells were grown in the presence or absence of 1.0 μg/ml C-CPE (*D* and *E*). Scale bars, 20 μm. Similar results were obtained from another set of experiments for *D* and *E*.

Notably, AKT and SGK1 were associated with transiently introduced ERα, but not with ERαΔC, in Ishikawa:*ESR1^−/−^* cells (*Figure 5A* and B), indicating that both kinases target either the LBD/AF2 domain (E region) or F region of ERα, We next determined whether the CLDN6 signaling directed to ERαS518 and ligand (estradiol)-independently stimulated the ERα activity in endometrial cancer cells, as in the human breast cancer cell line MCF-7 (19). To this end, we generated Ishikawa:*CLDN6*:*ESR1^−/−^*:*ESR1-wt (wild-type)* and Ishikawa:*CLDN6*:*ESR1^−/−^*:*ESR1S518A* cells, in the latter of which ERαS518 was substituted for an alanine residue, and performed rescue experiments. These cells were also grown in phenol red-free medium with charcoal-treated FBS to exclude fat-soluble ligands. As shown in *Figure 5C*, the level of ERα protein in Ishikawa:*CLDN6*:*ESR1^−/−^*:*ESR1-wt* cells was similar to that in Ishikawa:*CLDN6*:*ESR1^−/−^*:*ESR1S518A* cells. The transcript levels of the four ER target genes (*BCL2*, *CCND1, MYC,* and *VEGFA*; ref. 24) were significantly higher in Ishikawa:*CLDN6* cells than in Ishikawa cells (*Figure 5D*). More importantly, the expression levels of these target genes were significantly reduced in Ishikawa:*CLDN6*:*ESR1^−/−^*:*ESR1S518A* cells compared with those in Ishikawa:*CLDN6*:*ESR1^−/−^*:*ESR1-wt* cells (*Figure 5D*). Furthermore, cell proliferation was decreased in in Ishikawa:*CLDN6*:*ESR1^−/−^*:*ESR1S518A* and HEC-1A:*CLDN6*:*ESR1S518A* cells compared with those in Ishikawa:*CLDN6*:*ESR1^−/−^*:*ESR1-wt* and HEC-1A:*CLDN6*:*ESR1-wt* cells, respectively (*Figure 5E*). In addition, cell migration was significantly diminished in Ishikawa:*CLDN6*:*ESR1^−/−^*:*ESR1S518* cells compared with those in Ishikawa:*CLDN6*:*ESR1^−/−^*:*ESR1-wt* cells (*Figure 5F*). Hence, the CLDN6-adhesion signaling directed to ERαS518 for promoting the ERα activity and malignant phenotypes in endometrial cancer cells.

**Figure 5.**
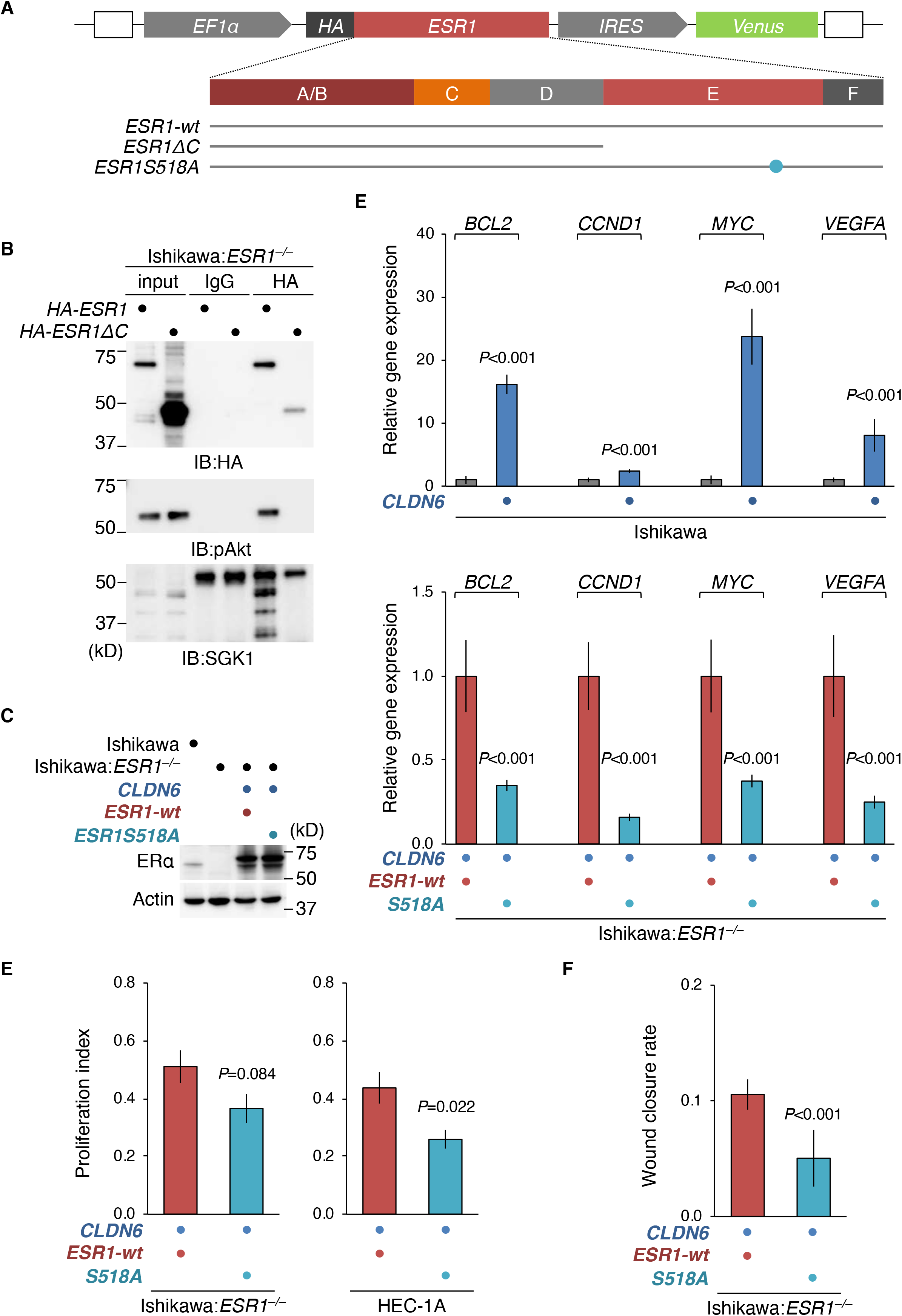
The CLDN6 signaling targets ERαS518 in endometrial carcinoma cells. (*A*) The construct of wild-type and mutant HA-ESR1 expression vectors. (*B*) Association of between either pAKT or SGK1 and ERα in Ishikawa:*ESR1^−/−^* cells transiently transfected with the HA-ESR1 expression vector. In the input lanes, 10% for HA, 1% for SGK1, or 0.1% for AKT of the input protein samples were loaded. (*C*) Western blot for the indicated proteins in the revealed Ishikawa cells. (*D*) RT-qPCR for the indicated molecules in the revealed Ishikawa cells. The expression levels of target genes are normalized to the corresponding *GAPDH* levels, and the relative levels are shown in the histograms (mean ± SD; *n* = 6) from two independent experiments. (*E*) BrdU assay for the indicated Ishikawa and HEC-1A cells. The BrdU/DAPI levels are shown in histograms (mean ± SD; *n* = 6). (*F*) Wound healing assay for the revealed Ishikawa cells. The values represent wound closure rates at day 2 (mean ± SD; *n* = 16). Similar results were obtained from another set of experiments for *D* and *E*.

### The CLDN6 signaling ERα-dependently and independently modulates gene expression in endometrial carcinoma cells

To identify downstream molecules that expression is altered by the CLDN6 signaling, we next compared, using RNA sequencing, the transcriptome in Ishikawa:*CLDN6* cells with that in Ishikawa cells (*Supplementary Table S5*). Following the analysis, we listed up the top 50 genes that expression was significantly down- or up-regulated in both of two distinct Ishikawa:*CLDN6* cell lines compared with parental Ishikawa cells (*Figure 6A*). Among such CLDN6-activated genes, we found the gene products associated with malignant phenotypes, including the soluble factors *CXCL1* (C-X-C motif ligand 1; ref. 25), *NRG1* (Neuregulin 1; ref. 26) and *NTN4* (Netrin 4; ref. 27), as well as the transmembrane receptor-associated tyrosine kinase *AXL* (28). We then by semi-quantitative RT-PCR clarified the expression of these four genes in Ishikawa, Ishikawa:*CLDN6*, Ishikawa:*ESR1^−/−^* and Ishikawa:*ESR1^−/−^*:*CLDN6* cells (*Figure 6B*). CLDN6 appeared to induce the expression of *AXL*, *NRG1* and *NTN4* transcripts via ERα. By contrast, CLDN6 activated the *CXCL1* mRNA expression in an ERα-independent manner. Thus, the CLDN6-activated genes can be classified into at least two groups with distinct ERα-dependence.

**Figure 6.**
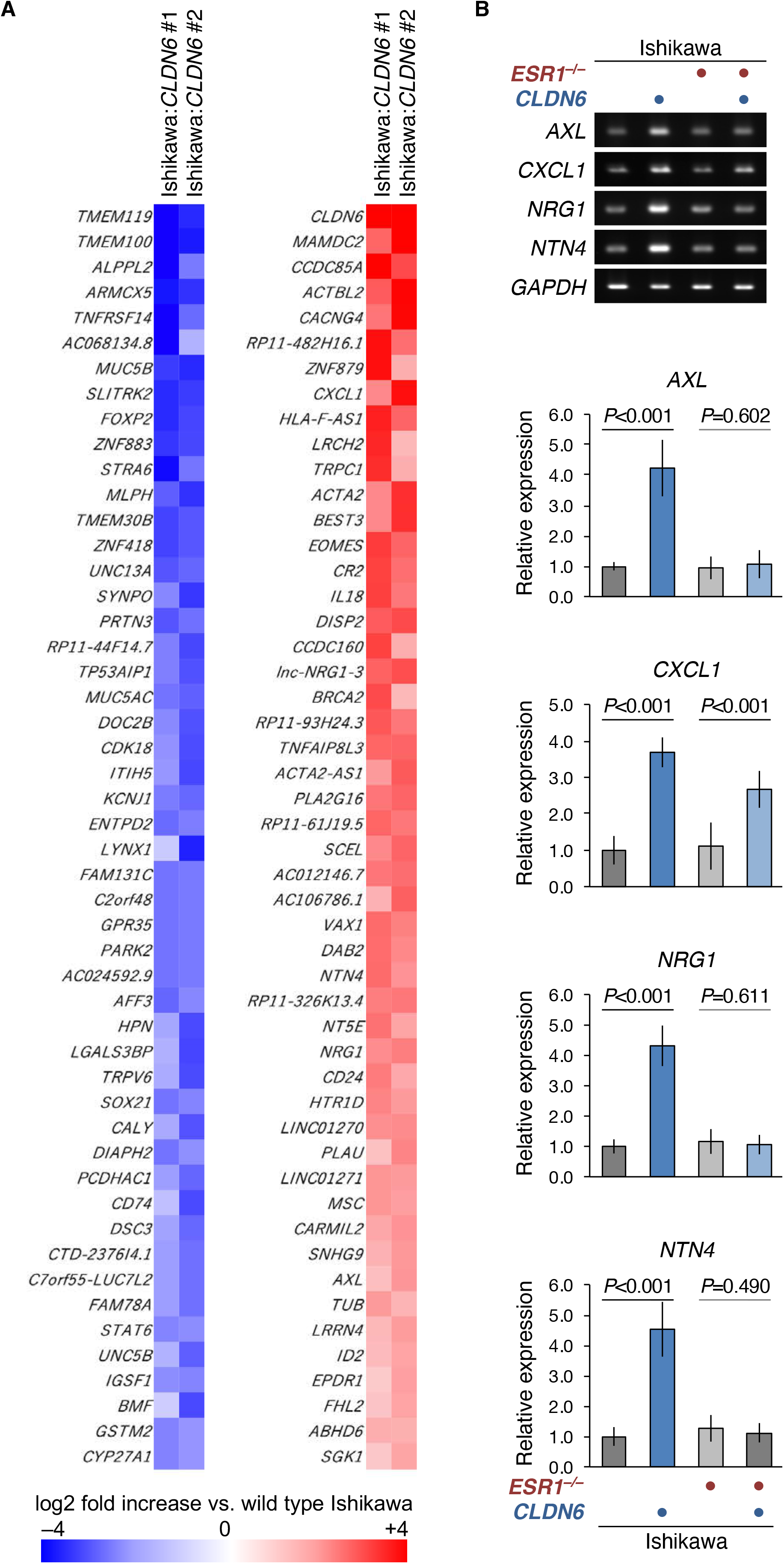
The CLDN6 signaling ERα-dependently and independently activates genes in endometrial carcinoma cells. (*A*) A heatmap of the top 50 genes that expression was significantly down- or up-regulated in both of two distinct Ishikawa:*CLDN6* cell lines compared with parental Ishikawa cells (*P*<0.05). (*B*) RT-PCR analysis for the indicated genes in the revealed cell lines. The expression levels relative to *GAPDH* are shown in the histograms (mean ± SD; *n* =7) from three independent experiments.

## Discussion

In the present study, we demonstrated that high CLDN6 expression in endometrial cancer tissues, in which the strong and moderate signal intensity on cell membranes was observed at greater than 30% and 50%, respectively, was significantly related to several clinicopathological features such as surgical stages III/IV, histological grade 3, LVSI, lymph node metastasis and distant metastasis. Importantly, the high CLDN6 expression represented an independent prognostic factor (HR 3.50), and the five-year survival rate was about 30%, which was one third of that in the low expression group. Thus, the aberrant CLDN6 expression appeared to corelate with poor outcome in patients with endometrial cancer. Taken together with the finding that CLDN6 is barely expressed in normal adult cells as described above, the established anti-human CLDN6 mAbs would provide powerful tools that selectively recognize CLDN6 protein in a range of cancer tissues.

CLDNs comprise a gene family as described above, and some anti-CLDN Abs are known to react not only with the corresponding CLDN but also with other CLDN subtypes (29). Therefore, it is of particular importance to verify the specificity of the anti-CLDN Abs used. Along this line, we previously established the anti-CLDN pAbs that selectively recognize CLDN1, CLDN5, CLDN6, CLDN7, CLDN12 or CLDN15 as far as we determined (22,30,31). The anti-CLDN6 pAb is one of the most reliable anti-CLDN6 Abs, and is used for immunohistochemical staining of formalin-fixed paraffin-embedded human tissues (18,32,33). However, we noticed in the present work that it also reacted with highly expressed CLDN4 and CLDN5 less efficiently than CLDN6, reinforcing the importance of validating the selectivity of each anti-CLDN Ab.

We also showed that CLDN6 accelerated endometrial cancer progression *in vitro* and *in vivo*. This was obvious because introduction of the human *CLDN6* gene was enough to promote cell proliferation and migration in two distinct endometrial cancer cell lines Ishikawa and HEC-1A:*ESR1*. In addition, overexpression of CLDN6 in Ishikawa cells led to enhanced tumor growth and invasion into the fibrous capsule in xenografts. Thus, we established the clinicopathological and biological relevance of the high CLDN6 expression in endometrial cancer.

Another finding of the present study is that the EC2 and Y196/200 of CLDN6 are responsible for recruiting and activating SFKs in endometrial cancer cells, as well as promoting the malignant properties. This conclusion was drawn from the following results: 1) the pSFK levels were increased in Ishikawa:*CLDN6* cells but not in Ishikawa:*CLDN6Y196A* or Ishikawa:*CLDN6Y200A* cells; 2) colocalization of CLDN6 and pSFK along cell boundaries was evident in Ishikawa:*CLDN6* cells, and diminished by C-CPE treatment; 3) a CLDN6-pSFK complex was formed in Ishikawa:*CLDN6* cells, and their association was decreased upon C-CPE exposure and in Ishikawa:*CLDN6Y196A* and Ishikawa:*CLDN6Y200A* cells; 4) the increased cell growth and migration in both Ishikawa:*CLDN6* and HEC-1A:*ESR1:CLDN6* cells were abrogated upon C-CPE treatment; 5) the CLDN6-stimulated cell proliferation was not detected in Ishikawa:*CLDN6Y196A* or Ishikawa:*CLDN6Y200A* cells. We also demonstrated that SFKs in turn phosphorylated CLDN6 at both Y196 and Y200, and tyrosine-phosphorylation of CLDN6 was governed by the EC2 domain. We previously reported that similar reciprocal regulation between CLDN6 and SFKs is also observed in mouse F9 stem cells (19), further strengthening our conclusion. Moreover, using the respective protein kinase inhibitors, we revealed that the PI3K-dependent AKT and SGK cascades contributed to the CLDN6/SFK signaling in endometrial cancer progression.

The most important conclusion of the present work is that the CLDN6/SFK/PI3K-dependent AKT and SGK signalings target ERα in endometrial cancer cells. This was apparent because CLDN6-accerelated cell growth and migration were hindered in Ishikawa:*CLDN6*:*ESR1^−/−^* cells. Using HEC-1A expressing CLDN6 and/or ERα, it was confirmed that the CLDN6 signaling in endometrial cancer advancement was mediated via ERα. Furthermore, AKT and SGK1 formed a complex with ERα in endometrial cancer cells, reinforcing the conclusion. On the other hand, neither kinases were associated with ERαΔC, indicating that they do not directly target the known AKT substrate S167 (34–37) at least in endometrial cancer cells. Instead, our RT-qPCR analysis indicated that the CLDN6 signaling directed to S518 in ERα and ligand-independently activated various oncogenic target genes. We also revealed that ERα-S518 is responsible for the CLDN6-accelerated malignant behaviors in endometrial cancer cells. The pathobiological relevance of the ERαS518 phosphorylation should be determined not only in endometrial cancer tissues, but also in other hormone-dependent tumors, such as ovarian cancer and breast cancer, in future experiments.

Our RNAseq analysis revealed that a variety of gene expression, including the *SGK1* gene, was altered between Ishikawa and Ishikawa:*CLDN6* cells. Among four representative gene products associated with tumor progression in various cancers, the *AXL*, *NRG1* and *NTN4* genes were activated by the ERα-dependent CLDN6 signaling. By contrast, the expression of *CXCL1* transcripts was induced by CLDN6 in an ERα-independent manner. Enhanced cell migration was observed in HEC-1A:*CLDN6* cells without ERα expression, supporting the presence of not only ERα-dependent but also ERα-independent CLDN6 signaling. Interestingly, the novel AKT/SGK-consensus phosphorylation motif is conserved in 14 of 48 members of human nuclear receptors (19). Taken together, CLDN6 may also target these nuclear receptors and possibly other transcription factors in order to regulate the expression of certain genes. In fact, we previously reported that the CLDN6 signal targets RARγ in mouse F9 stem cells to initiate epithelial differentiation (19).

Genomic and non-genomic heterogeneity among distinct cell populations within cancers is known to influence tumour behaviors (38). Our immunohistochemical study revealed intratumor heterogeneity on the CLDN6 expression within human endometrial cancer and Ishikawa:*CLDN6* xenograft tissues. These tumors were composed of CLDN6-positive and negative subpopulations, even in endometrial cancer tissues with high CLDN6 expression. Hence, the expression of CLDN6 should be carefully evaluated when small biopsy specimens and tissue arrays were subjected to immunohistochemistry. Of note, since the gene expression of various diffusive factors was induced in Ishikawa:*CLDN6* as described above, non-cell-autonomous paracrine effects between CLDN6-positive and negative cancer cells may also contribute to the enhancement of tumor progression.

In summary, we here established that high expression of CLDN6 protein in endometrial cancer leads to more aggressive tumors and predicts poor prognosis. We also demonstrated that the CLDN6/SFK/PI3K-dependent AKT and SGK cascades direct to S518 in human ERα and stimulated its activity, resulting in progression of tumor behaviors in endometrial cancer. Therefore, in addition to the PI3K/AKT pathway, which is frequently altered in endometrial cancers (39–42), the CLDN6/SFK, SGK and ERαS518 may be promising therapeutic targets for endometrial cancer. It would also be interesting to determine whether a similar link between cell adhesion and nuclear receptor signaling regulates tumor progression in various types of cancers.

## Materials and Methods

### Antibodies

The antibodies used in this study are listed in *Supplementary Table S1*. A rabbit pAb against CLDN6 was generated in cooperation with Immuno-Biological Laboratories as described previously (<u>22</u>).

Rat mAbs against CLDN6 were established using the iliac lymph node method (<u>21</u>). In brief, a polypeptide, (C)SRGPSEYPTKNYV corresponding to the cytoplasmic domain of CLDN6, was coupled via the cysteine to Imject™ Maleimide-Activated mcKLH (Thermo Fisher SCIENTIFIC). The conjugated peptide was intracutaneously injected with Imject™ Freund’s Complete Adjuvant (Themo Fisher SCIENTIFIC) into the footpads of anesthetized eight-week-old female rats. The animals were sacrificed 14 days after immunization, and the median iliac lymph nodes were collected, followed by extraction of lymphocytes by mincing. Extracted lymphocytes were fused with cells of the SP2 mouse myeloma cell line by polyethylene glycol. Hybridoma clones were maintained in GIT medium (Wako) with supplementation of 10% BM-Condimed (Sigma-Aldrich). The supernatants were screened by ELISA.

### Tissue collection, immunostaining, and analysis

Paraffin-embedded tissue sections were obtained from 173 patients with uterine endometrial cancer who underwent hysterectomy, bilateral saplingo-oophrectomy, and/or lymphadenectomy between 2003 and 2012 at Fukushima Medical University Hospital (FMUH) and Iwaki City Medical Center (ICMC). Informed consent was obtained from all the patients. The subjects were limited to patients with confirmed 5-year outcomes and who died due to uterine endometrial cancer and metastasis. The clinicopathological characteristics of patients are summarized in *Supplementary Table S2*. The detailed information, including postoperative pathology diagnosis reports, age, stage (FIGO2008), histological type, histological grade, lymph-vascular space invasion (LVSI), lymph node metastasis, distant metastasis, overall survival (OS), and recurrence-free survival (RFS), were also obtained. The staging of patients between 2003 and 2007 were modified in accordance with the FIGO 2008 systems. Distant metastasis was judged by diagnostic imaging. The study was approved by the ethics committee of FMUH and ICMC.

For immunostaining, uterine endometrial cancer tissues were obtained, and the 10% formalin-fixed and paraffin-embedded tissue blocks were sliced into 5-μm-thick sections, then deparaffinized with xylene and rehydrated using a graduated series of ethanol. The sections were immersed in 0.3% hydrogen peroxide in methanol for 20 min at room temperature to block endogenous peroxidase activity. Antigen retrieval was performed by incubating the sections in boiling citric acid buffer (pH 6.0) for antigen retrieval in a microwave. After blocking with 5% skimmed milk at room temperature for 30 min, the sections were incubated overnight at 4°C with the primary antibodies. Histofine SAB-PO kit for rabbit (Nichirei) or VECTASTAIN Elite ABC HRP Kit for rat (VECTOR LABORATORIES) was used for 3’,3’-diaminobenzidine (DAB) staining.

Immunostaining results were interpreted by three independent pathologists and one gynecologist using a semi-quantitative scoring system, immunoreactivity score (IRS; 46). The immunostaining reactions were evaluated according to signal intensity (SI: 0, no stain; 1, weak; 2, moderate; 3, strong) and percentage of positive cells (PP: 0, <1%; 1, 1 to 10%; 2, 11 to 30%; 3, 31 to 50%; and 4, >50%). The SI and PP were then multiplied to generate the IRS for each case. To determine the optical cut-off values of IRS for CLDN6 expression, the receiver operating characteristics (ROC) curve was plotted and analysed. Based on this analysis, we divided the samples into two groups based on the results of the immunostaining in the tissues: low expression (IRS<8) and high expression (IRS2:8; *Supplementary Table S3*).

### Cell lines and cell culture

The Ishikawa cell line was obtained from Kasumigaura Medical Center and from Dr.Yamada (Wakayama Medical University). The HEC-1A cell line (<u>43</u>) were obtained from National Institute of Biomedical Innovation, Health and Nutrition (Japan). F9:*Cldn6* was previously established (<u>16</u>). Cells were grown in Roswell Park Memorial Institute (RPMI) 1640 (Ishikawa), McCoy’s 5A (HEC-1A), or Dulbecco’s Modified Eagle Medium (DMEM; HEK293T), with 10% Fetal bovine serum (FBS; Sigma-Aldrich) and 1% Penicillin-streptomycin mixture (Gibco, Waltham, MA). Ishikawa and HEC-1A cells were treated with 1 μM of C-CPE, 10 μM of PP2 (Calbiochem), 10 μM of LY294002 (Cell Signaling TECHNOLOGY), 10 μM of AKT inhibitor VIII (funakoshi), or 0.1 nM of SGK-1 inhibitor (Santa Cruz Technology) one or two days after plating. For preparation of charcoal-treated FBS, 500 ml of FBS was treated with 0.5 g of Charcoal, dextran coated (Sigma) overnight at 4°C followed by filtration using 0.22 μm cellulose acetate filter membranes. C-CPE production and purification were performed as described previously by using E.coli BL21 and the expression vector pET16b coding C-CPE194-319 (<u>19</u>).

### Expression vectors, transfection, and establishment of stable cell lines

The protein coding regions of human *CLDN1*, *CLDN4*, *CLDN5*, *CLDN6*, *CLDN9*, and *ESR1* were cloned into the *BamH*I/*Not*I site of the CSII-EF-MCS-IRES2-Venus (RIKEN, RDB04384) plasmid. Hemagglutinin (HA) tag was added by PCR with tailed primer. Expression vectors of mutant genes (*CLDN6Y196A*, *CLDN6Y200A*, *ESR1ΔC*, and *ESR1S518A*) were established by a standard site-directed mutagenesis protocol using KOD -Plus-Mutagenesis Kit (TOYOBO) following the providers protocol.

For transient expression of the target genes (*CLDN1*, *CLDN4*, *CLDN5*, *CLDN6*, *CLDN9*; *Supplementary Figure S1*), 5×10^6^ cells were transfected with 10 μg of the indicated vectors using 30 μg of Polyethylenimine Max (PEI Max, Cosmo Bio) 8 h after passage.

Ishikawa:*CLDN6*, Ishikawa:*CLDN6Y196A*, Ishikawa:*CLDN6Y200A*, Ishikawa:*ESR1^−/−^:ESR1*, Ishikawa:*ESR1^−/−^:ESR1ΔC*, Ishikawa:*ESR1^−/−^:CLDN6:ESR1S518A*, HEC1A:*CLDN6*, HEC1A:*ESR1*, and HEC1A:*ESR1:CLDN6* cell lines were established by lentiviral transfection. First, lentiviral vectors (*CLDN6*, *CLDN6Y196A*, *CLDN6Y200A*, *ESR1*, *ESR1ΔC*, and *ESR1S518A*) were generated by transfecting HEK293T cells with 10 μg of the CSII plasmids containing the target genes, 5 μg of packaging plasmids psPAX2 (Addgene, #12260) and pCMV-VSV-G (Addgene, #8454) using PEI Max. Culture media containing recombinant lentiviruses were collected 72 h after transfection. The lentiviral vectors were added to cell culture medium of Ishikawa, Ishikawa:*ESR1^−/−^*, or HEC-1A cell lines after filtration. More than 48 h after transfection, the cells were used for further analysis. Ishikawa:*CLDN6* cell lines were single cell-cloned by limiting dilution with 96-well culture plates.

### Genome editing

To establish the Ishikawa:*ESR1^−/−^* cell line, a pair of transcription activator-like effector nucleases (TALENs) targeting the 2nd exon of *ESR1* gene were designed by TALEN Targeter 2.0 software (https://tale-nt.cac.cornell.edu/node/add/talen; <u>ref.</u> <u>44</u>). The expression vector of the TALENs were cloned by using Platinum TALEN Kit (<u>45</u>). The plasmids were transiently transfected by Polyethylenimine Max (PEI Max, Cosmo Bio). Next, 24-48 h after transfection, the cells were exposed to 100 μg/ml of hygromycin for positive selection, followed by limiting dilution and genotyping by PCR-based restriction fragment length polymorphism (RFLP; <u>ref.</u> <u>46</u>) utilizing the endogenous *Stu* I recognition site. Knockout of *ESR1* genes was verified by DNA sequence after TA-cloning of genomic PCR products.

### Immunoprecipitation and immunoblot

Immunoprecipitation was performed using an Immunoprecipitation kit (Protein G, Sigma), following the manufacturer’s protocol. Immunoblot analysis was performed as previously described (47). Each blot was stripped with Restore Western blot stripping buffer (Pierce Chemical) and immunoprobed with anti-actin antibody. Signals in the immunoblots were quantified using ImageJ software (Wayne Rasband National Institutes of Health). The protein levels were normalized to the corresponding actin levels, and their relative levels were then presented.

### RNA extraction, RT-PCR, and RNA sequencing

For analysis of gene expression, total RNA was isolated from cells using TRIzol RNA Isolation Reagents (Thermo Fisher Scientific), and reverse transcription was performed using Primescript II RT Kit (Clontech). For PCR, the target sequences were amplified using GoTaq Green Master Mix (Promega). The primers for RT-PCR are listed in *Supplementary Table S4*. Aliquots of the PCR products were loaded onto 2.5% agarose gel and analyzed after staining with ethidium bromide. Original PCR values were quantified by ImageJ software (Wayne Rasband National Institutes of Health). The expression levels of the target genes in RT-PCR were divided by the corresponding *GAPDH* signal intensity. Quantitative PCR (qPCR) was performed using THUNDERBIRD SYBR qPCR Mix (TOYOBO) and Step One Real-Time PCR System (Applied Biosystems).

RNA sequencing and mapping were performed by TaKaRa Bio Inc. For mapping, the index trimmed single-end 100 bp reads were aligned to the human reference genome (GRCh38 v90) to generate bam files. The mapped bam files were imported to SeqMonk software (Babraham Bioinformatics; https://www.bioinformatics.babraham.ac.uk/projects/seqmonk/) as single ended RNA-Seq data. Then they were quantitated by using the default RNA-Seq quantitation pipline. *P*-value was calculated from the intensity difference and representative genes of showing significant change (*P*<0.05) are listed in Figure 6A. The normalized expression levels of each gene are shown in *Supplementary Table S5*.

### Fluorescence Immunohistochemistry

Cells were grown on coverslips coated by Cellmatrix Type I-A (Nitta gelatin). The samples were fixed in 1% paraformaldehyde and 0.1% Triton-X for 10 min at room temperature. After washing with PBS, they were preincubated in PBS containing 5% skimmed milk. They were subsequently incubated overnight at 4°C with primary antibodies in PBS, then rinsed again with PBS, followed by a reaction for 1 h at room temperature with appropriate secondary antibodies. All samples were examined using a laser-scanning confocal microscope (FV1000, Olympus). Photographs were processed with Photoshop CC (Adobe) and ImageJ software (Wayne Rasband National Institutes of Health).

### Cell proliferation, migration, and apoptosis assays

Cell proliferation index was evaluated by incorporation of bromodeoxy uridine (5-Bromo-2-DeoxyUridine, BrdU, sigma). Cells were exposed to BrdU for 5 min after passage. The specimens were fixed with 4% paraformaldehyde and 0.1% Triton-X, followed by immunostaining with anti-BrdU antibody (BD) and its standard protocol.

For evaluating cell migration, wound areas were generated by scratching with disposable 1,000 μl pippette tips 24-48 h after passage. Culture media were changed daily. Photographs of the wound areas were taken at the same locations, using a phase-contrast microscope. Wound healing was calculated as the percentage of the remaining cell-free area compared with the initial wound area using ImageJ software.

*in situ* Cell Death Detection Kit (Roche) was used for evaluation of cell apoptosis.

### Xenograft model

Xenograft studies were performed in 8-week-old NOD/ShiJic-scid female mice (CLEA-Japan). 1 × 10^7^ cells were subcutaneously injected into the back of anesthetized mice. Then, 28 d after injection, the mice were ethically sacrificed. All animal experiments conformed to the National Health Guide for the Care and Use of Laboratory Animals, and were approved by the Animal Committee at Fukushima Medical University.

### Statistical analysis

We used the chi-squared test to evaluate the relationship between CLDN6 expression and various clinicopathological parameters (age, stage, histological type, histological grade, LVSI, lymph node metastasis, distant metastasis, 5-year OS, and 5-year RFS). Survival analysis was performed using the Kaplan-Meier method, and differences between the groups were analyzed using the log-rank test. The Cox regression multivariate model was used to detect the independent predictors of survival. Two-tailed P-values less than 0.05 were considered to indicate a statistically significant result. All statistical analyses were performed using SPSS software version 23.0 (IBM).

Statistical significance for cell proliferation, migration, xenograft studies, as well as for semi quantitative and quantitative PCR analyses, was analyzed by unpaired two-tailed t-test.

## Acknowledgements

We thank A. Hozumi and K. Watari for their technical assistance; and English Editing Service of the Medical Research Promotion Office, Fukushima Medical University for their assistance with the manuscript.

## Supplementary Figure legends

**Figure S1.**
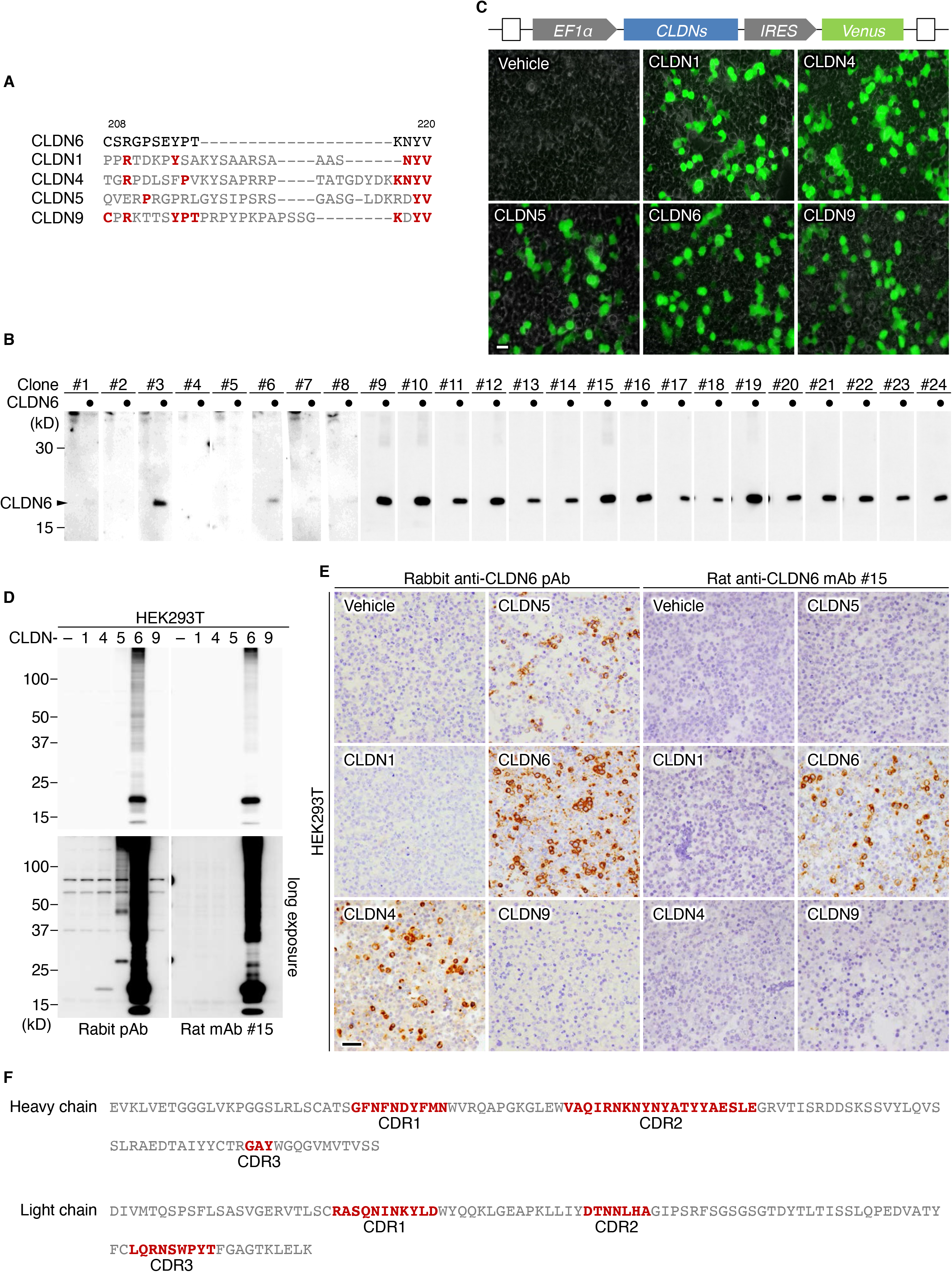
Generation of rat anti-human CLDN6 mAbs. (*A*) Amino acid sequences of the antigenic peptide of the C-terminal cytoplasmic domains of human CLDN6 and the corresponding regions of the closely related CLDNs. Conserved amino acids are shown in red. (*B*) The construct of CLDN1/4/5/6/9 expression vectors and the representative fluorescence images of the transfected HEK293T cells. EF-1α, elongation factor-1α; IRES, internal ribosome entry site. (*C*) Twenty-four hybridoma clones were screened by Western blot for CLDN6 in HEK293T cells that transiently transfected with the CLDN6 or empty expression vector. (*D* and *E*) HEK293T cells were transfected with individual CLDN expression vectors, and subjected to Western blot and imnunohistochemical analyses using the indicated anti-CLDN6 Abs. (*F*) The complementary determining-regions (CDRs) of an anti-human CLDN6 mAb (clone #15). Scale bars, 100 μm.

**Figure S2.**
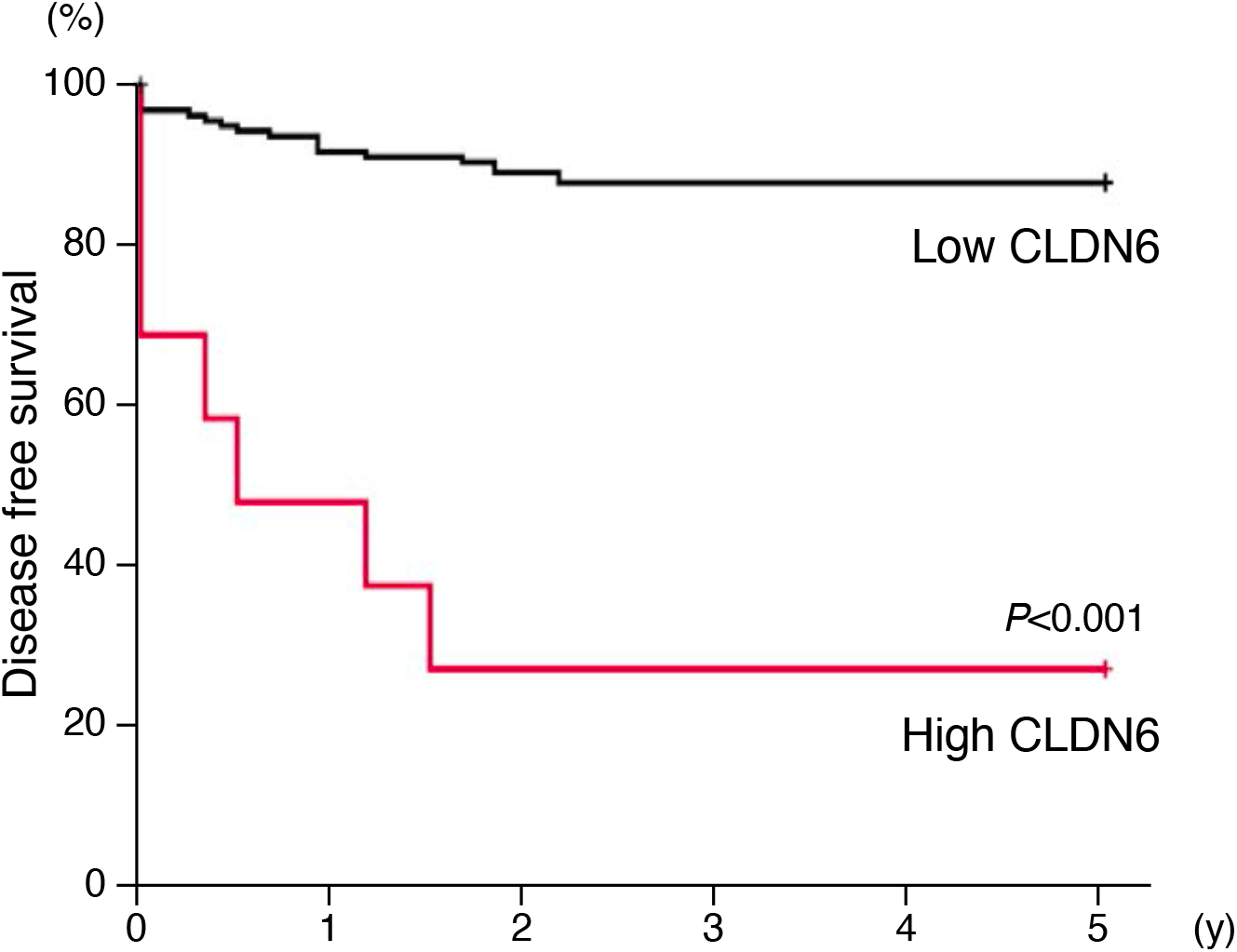
The 5-year recurrence-free survival for high and low CLDN6 expression groups in endometrial cancer subjects.

**Figure S3.**
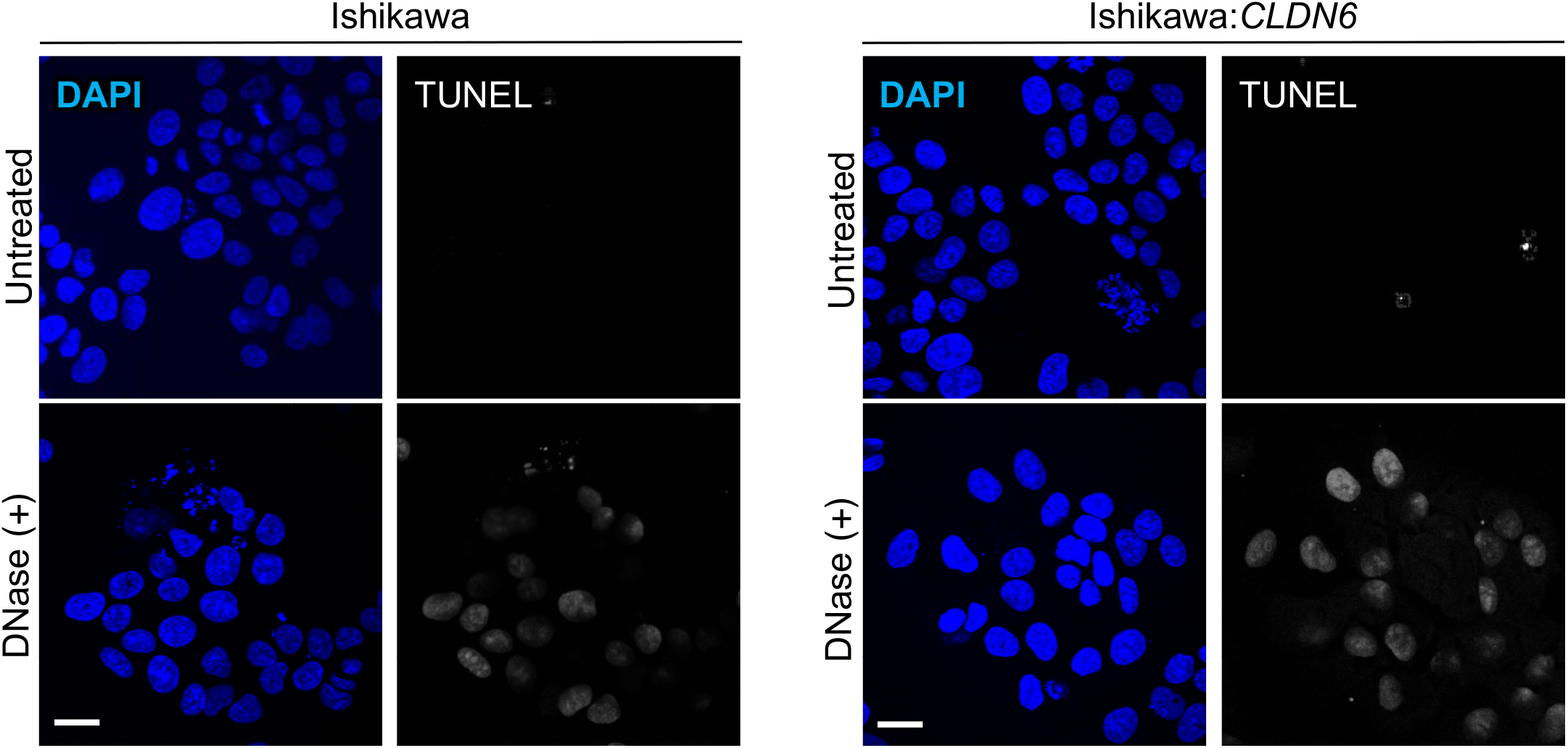
Apoptosis are not detected in Ishikawa and Ishikawa: *CLDN6* cells. Cells are subjected to TUNEL assay together with DAPI staining. As a positive control, cells were treated with DNase. Scale bars, 20 μm. Similar results were obtained from three independent experiments.

**Figure S4.**
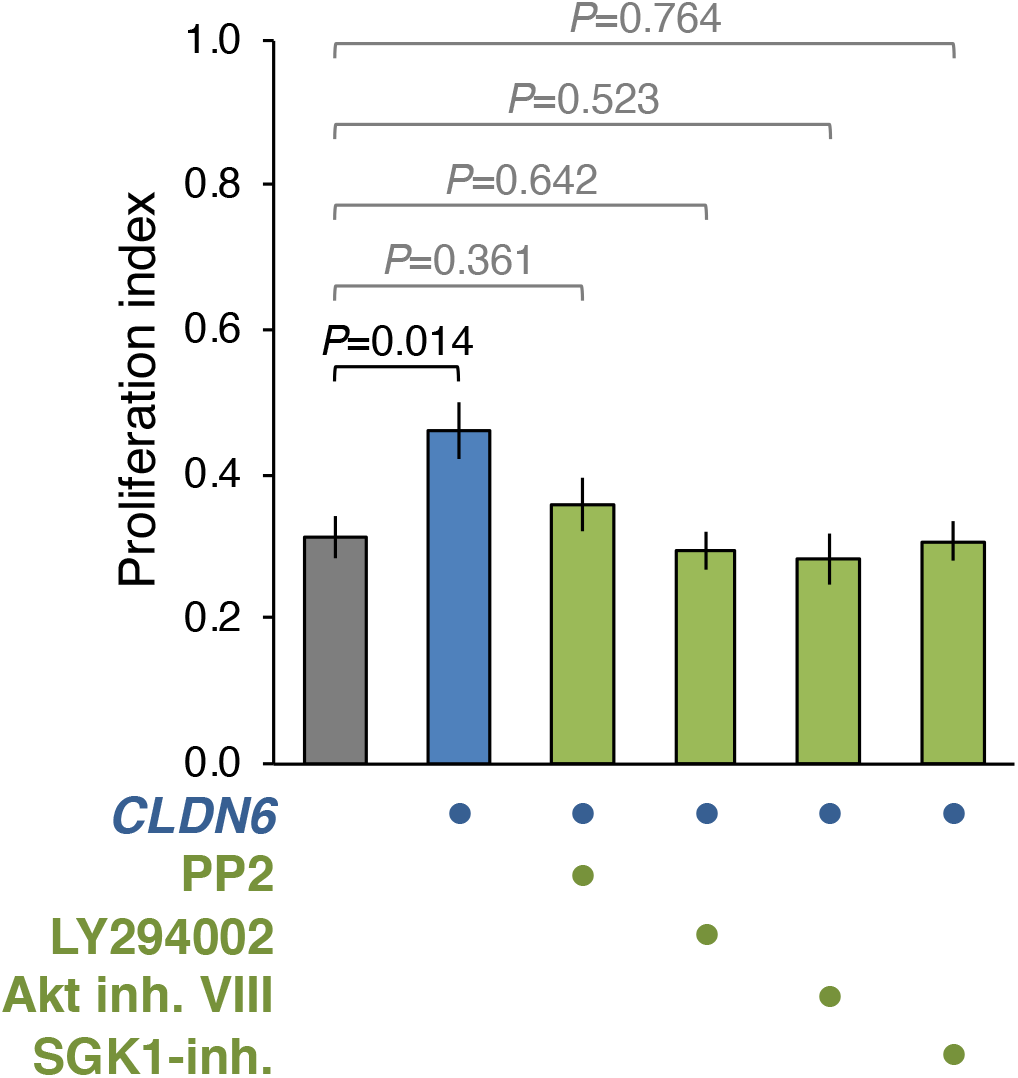
The SFK/PI3K-dependent AKT and SGK pathways are involved in the CLDN6-accererated endometrial cancer proliferation. Ishikawa and Ishikawa:*CLDN6* cells were grown in the presence of the vehicle, SFK, PI3K, AKT and SGK1 inhibitors (PP2, LY294002 and AKT inh VIII, 10 μM; SGK1 inh, 1 nM). The BrdU/DAPI levels are shown in histograms (mean ± SD; *n* = 6). Similar results were obtained from another independent experiments.

**Figure S5.**
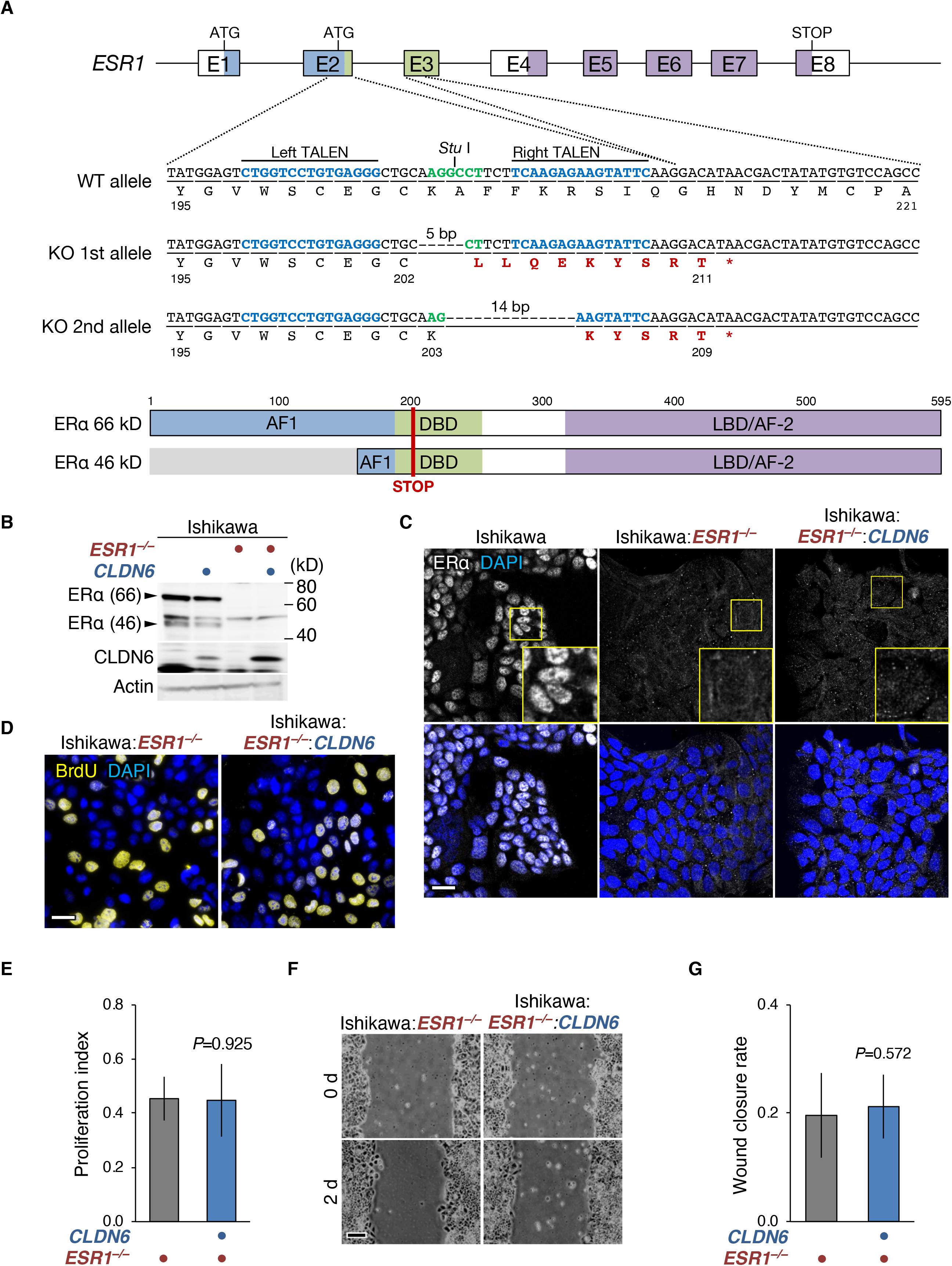
ERα is required for CLDN6-stimulated malignant phenotypes of endometrial carcinoma cells. (*A*) Knockout (KO) of the *ESR1* gene encoding human ERα in Ishikawa cells using the TALEN method. The KO in Ishikawa:*ESR1^−/−^* cells is confirmed by DNA sequencing. (*B* and *C*) Absence of ERα protein in Ishikawa:*ESR1^−/−^* and Ishikawa:*CLDN6*:*ESR1^−/−^* cells on Western blot (*B*) and immunofluorescence (*C*) analyses. (*D* and *E*) Representative (*D*) and quantitative (*E*) BrdU assay for the indicated cells. The BrdU/DAPI levels are shown in histograms (mean ± SD; *n* = 6). (*F* and *G*) Typical (*F*) and quantitative (*G*) wound healing assay for the indicated cells. The values represent wound closure rates (mean ± SD; *n* = 12). Scale bars, 50 μm (*F*); 20 μm (*C* and *D*). N.S., not significant. Similar results were obtained from another set of experiments for *E* and *G*.

**Table. S1.**

| antibodies | host | source | identifier | IP | IB | IHC |
| --- | --- | --- | --- | --- | --- | --- |
| AKT | rabbit | Cell Signaling Technology | 4691S |  | 1:1,000 |  |
| βActin | mouse | Thermo Fisher Scientific | A5441 |  | 1:10,000 |  |
| BrdU | rat | Creative Diagnostics | DPATB-H81231 |  |  | 1:1,000 |
| Claudin-6 | rabbit | Immuno-Biological Laboratories | 18865 | 2 μg | 1:2,000 | 1:200 |
| ERα(HC-20) | rabbit | Santa Cruz Technology | sc-543 |  | 1:1,000 | 1:200 |
| HA | rat | Roche | 11867423001 | 1 μg | 1:1,000 |  |
| Ki67 | mouse | Dako | M7240 |  |  | 1:200 |
| PI3K (55) | rabbit | Cell Signaling Technology | 11889S |  | 1:1,000 |  |
| Phospho-AKT | rabbit | Cell Signaling Technology | 4060S |  | 1:1,000 |  |
| Phospho-PI3K (85/55) | rabbit | Bioss | bs-3332R |  | 1:1,000 |  |
| Phospho-Tyrosine | mouse | Santa Cruz Technology | Sc-508 | 2 μg | 1:1,000 | 1:100 |
| Phospho-SFK (Tyr416) | rabbit | Cell Signaling Technology | 2101 |  | 1:1,000 | 1:100 |
| Phospho-SGK1 |  | MERK |  |  |  |  |
| SFK | mouse | Cell Signaling Technology | 2102S |  | 1:1,000 |  |
| SGK1 | rabbit | Cell Signaling Technology | 12103S |  | 1:1,000 |  |
| mouse IgG (HRP) | sheep | GE Health Care | NA931V |  | 1:10,000 |  |
| rabbit IgG (HRP) | donkey | GE Health Care | NA934V |  | 1:2,000 |  |
| rat IgG (HRP) | goat | GE Health Care | NA935V |  | 1:2,000 |  |
| rabbit IgG (Cy3) | donkey | Jackson ImmunoResearch | 711-165-152 |  |  | 1:300 |
| rat IgG (Cy3) | donkey | Jackson ImmunoResearch | 712-165-150 |  |  | 1:300 |
| rat IgG(Alexa Fluor 647) | donkey | Jackson ImmunoResearch | 712-605-153 |  |  | 1:300 |

IP, Immunoprecipitation; IB, Immunoblotting; IHC, Immunohistochemistry

**Table. S2.**
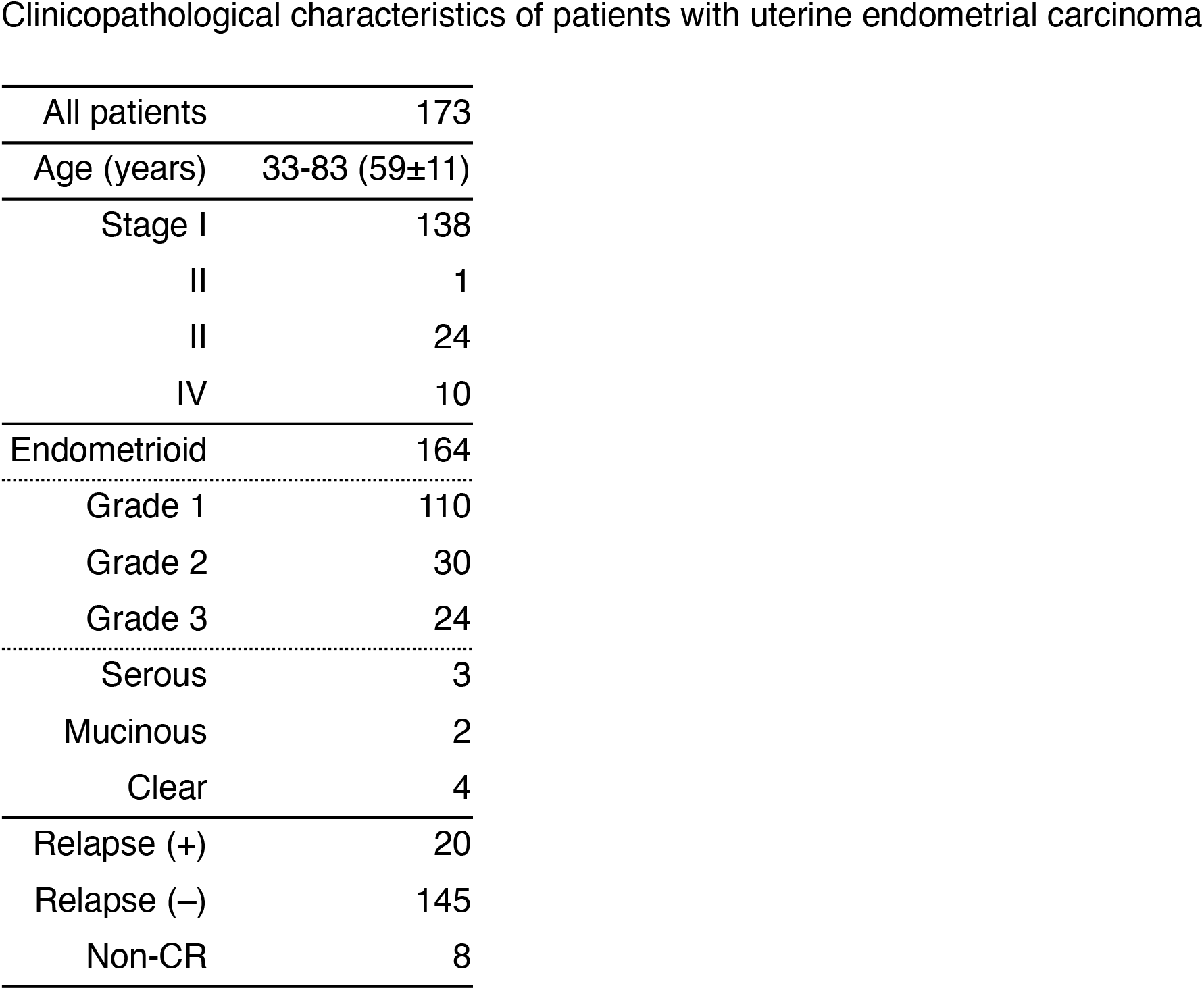

**Table. S3.** Immunoreactivity score (IRS)

| Score | Signal intensity (SI) | Percentage of positive cells (PP) |
| --- | --- | --- |
| 0 | negative | <1% |
| 1 | weak | 1-10% |
| 2 | moderate | 11-30% |
| 3 | strong | 31-50% |
| 4 |  | >50% |

| SI × PP | IRS |  |
| --- | --- | --- |
| 0 | Score 0 | CLDN6 low |
| 1–2 | Score 1+ |  |
| 3–6 | Score 2+ |  |
| 8–12 | Score 3+ | CLDN6 high |

**Table. S4.**
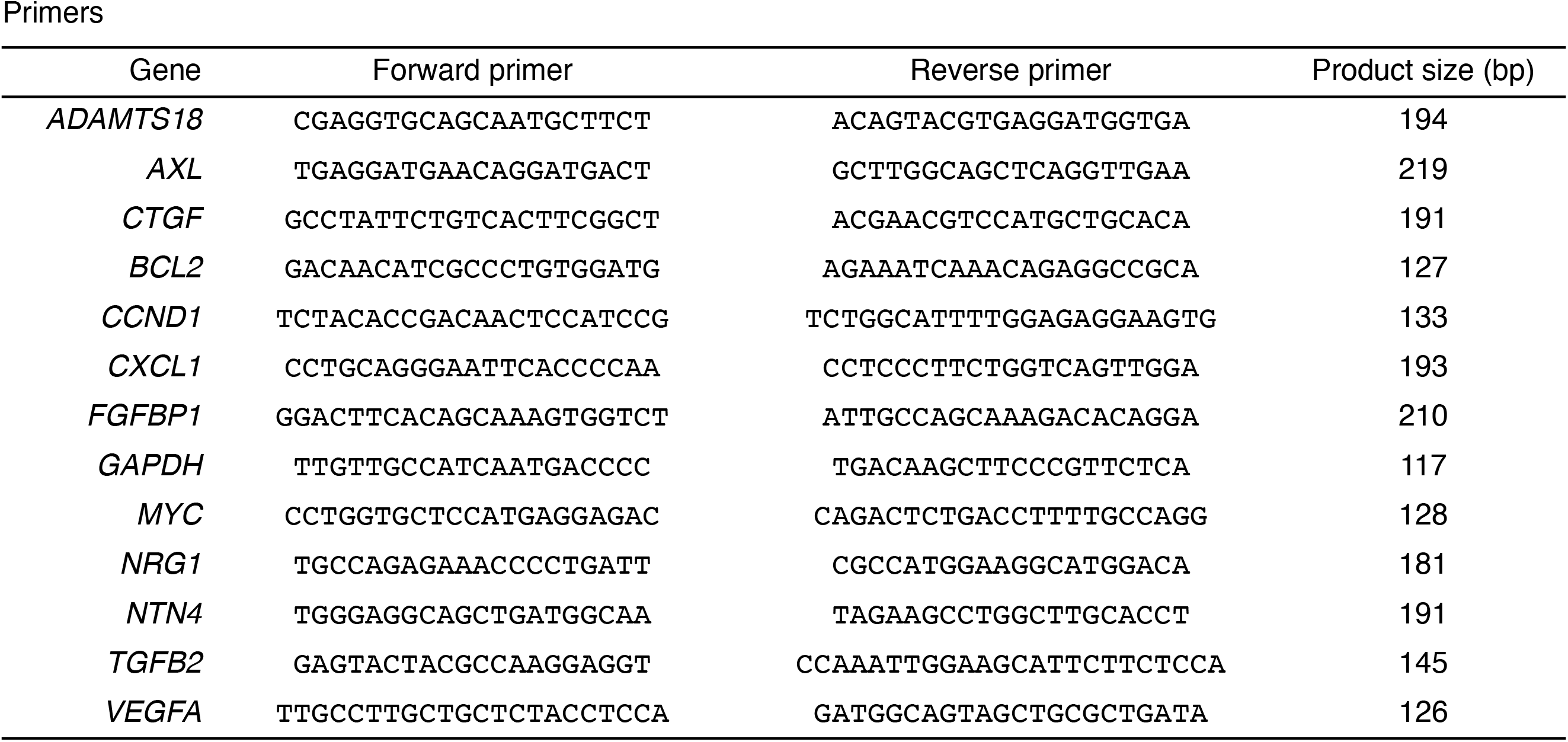

## References

1. Ferlay J, Steliarova-Foucher E, Lortet-Tieulent J, Rosso S, Coebergh JWW, Comber H, et al. Cancer incidence and mortality patterns in Europe: estimates for 40 countries in 2012. Eur J Cancer. 2013 Apr;49(6):1374–403.

2. Evans-Metcalf ER, Brooks SE, Reale FR, Baker SP. Profile of women 45 years of age and younger with endometrial cancer. Obstet Gynecol. 1998 Mar;91(3):349–54.

3. Kaaks R, Lukanova A, Kurzer MS. Obesity, endogenous hormones, and endometrial cancer risk: a synthetic review. Cancer Epidemiol Biomarkers Prev. 2002 Dec;11(12):1531–43.

4. Morice P, Leary A, Creutzberg C, Abu-Rustum N, Darai E. Endometrial cancer. Lancet. 2016 Mar 12;387(10023):1094–108.

5. Creasman WT, Heintz APM. Carcinoma of the corpus uteri. FIGO 26th Annual Report on the Results of Treatment in Gynecological Cancer. Int J Gynaecol Obstet. 2006 Nov;95 Suppl 1:S105–43.

6. Furuse M, Tsukita S. Claudins in occluding junctions of humans and flies. Trends Cell Biol. 2006 Apr;16(4):181–8.

7. Van Itallie CM, Anderson JM. Claudins and epithelial paracellular transport. Annu Rev Physiol. 2006;68:403–29.

8. Chiba H, Osanai M, Murata M, Kojima T, Sawada N. Transmembrane proteins of tight junctions. Biochim Biophys Acta. 2008 Mar;1778(3):588–600.

9. Zihni C, Mills C, Matter K, Balda MS. Tight junctions: from simple barriers to multifunctional molecular gates. Nat Rev Mol Cell Biol. 2016 Sep;17(9):564–80.

10. Tabariès S, Siegel PM. The role of claudins in cancer metastasis. Oncogene. 2017 Mar 2;36(9):1176–90.

11. Cavallaro U, Dejana E. Adhesion molecule signalling: not always a sticky business. Nat Rev Mol Cell Biol. 2011 Mar;12(3):189–97.

12. Kubota H, Chiba H, Takakuwa Y, Osanai M, Tobioka H, Kohama G, et al. Retinoid X receptor alpha and retinoic acid receptor gamma mediate expression of genes encoding tight-junction proteins and barrier function in F9 cells during visceral endodermal differentiation. Exp Cell Res. 2001 Feb 1;263(1):163–72.

13. Turksen K, Troy TC. Claudin-6: a novel tight junction molecule is developmentally regulated in mouse embryonic epithelium. Dev Dyn. 2001 Oct;222(2):292–300.

14. Chiba H, Gotoh T, Satohisa S, Kikuchi K, Osanai M, Sawada N. Hepatocyte nuclear factor (HNF)-4alpha triggers formation of functional tight junctions and establishment of polarized epithelial morphology in F9 embryonal carcinoma cells. Exp Cell Res. 2003 Jun 10;286(2):288–97.

15. Anderson WJ, Zhou Q, Alcalde V, Kaneko OF, Blank LJ, Sherwood RI, et al. Genetic targeting of the endoderm with claudin-6CreER. Dev Dyn. 2008 Feb;237(2):504–12.

16. Sugimoto K, Ichikawa-Tomikawa N, Satohisa S, Akashi Y, Kanai R, Saito T, et al. The tight-junction protein claudin-6 induces epithelial differentiation from mouse F9 and embryonic stem cells. PLoS ONE. 2013;8(10):e75106.

17. Sullivan LM, Yankovich T, Le P, Martinez D, Santi M, Biegel JA, et al. Claudin-6 is a nonspecific marker for malignant rhabdoid and other pediatric tumors. Am J Surg Pathol. 2012 Jan;36(1):73–80.

18. Ushiku T, Shinozaki-Ushiku A, Maeda D, Morita S, Fukayama M. Distinct expression pattern of claudin-6, a primitive phenotypic tight junction molecule, in germ cell tumours and visceral carcinomas. Histopathology. 2012 Dec;61(6):1043–56.

19. Sugimoto K, Ichikawa-Tomikawa N, Kashiwagi K, Endo C, Tanaka S, Sawada N, et al. Cell adhesion signals regulate the nuclear receptor activity. Proc Natl Acad Sci USA. 2019 Dec 3;116(49):24600–9.

20. Rodriguez AC, Blanchard Z, Gertz J. Estrogen Signaling in Endometrial Cancer: a Key Oncogenic Pathway with Several Open Questions. Horm Cancer. 2019 Jun;10(2-3):51–63.

21. Kishiro Y, Kagawa M, Naito I, Sado Y. A novel method of preparing rat-monoclonal antibody-producing hybridomas by using rat medial iliac lymph node cells. Cell Struct Funct. Japan Society for Cell Biology; 1995 Apr;20(2):151–6.

22. Satohisa S, Chiba H, Osanai M, Ohno S, Kojima T, Saito T, et al. Behavior of tight-junction, adherens-junction and cell polarity proteins during HNF-4alpha-induced epithelial polarization. Exp Cell Res. 2005 Oct 15;310(1):66–78.

23. Lang F, Böhmer C, Palmada M, Seebohm G, Strutz-Seebohm N, Vallon V. (Patho)physiological significance of the serum- and glucocorticoid-inducible kinase isoforms. Physiol Rev. 2006 Oct;86(4):1151–78.

24. Ikeda K, Horie-Inoue K, Inoue S. Identification of estrogen-responsive genes based on the DNA binding properties of estrogen receptors using high-throughput sequencing technology. Acta Pharmacol Sin. 2015 Jan;36(1):24–31.

25. Zhang Z, Chen Y, Jiang Y, Luo Y, Zhang H, Zhan Y. Prognostic and clinicopathological significance of CXCL1 in cancers: a systematic review and meta-analysis. Cancer Biol Ther. Taylor & Francis; 2019 Oct 10;20(11):1380–8.

26. Sheng Q, Liu X, Fleming E, Yuan K, Piao H, Chen J, et al. An activated ErbB3/NRG1 autocrine loop supports in vivo proliferation in ovarian cancer cells. Cancer Cell. 2010 Mar 16;17(3):298–310.

27. Hu Y, Ylivinkka I, Chen P, Li L, Hautaniemi S, Nyman TA, et al. Netrin-4 promotes glioblastoma cell proliferation through integrin β4 signaling. Neoplasia. 2012 Mar;14(3):219–27.

28. Rankin EB, Giaccia AJ. The Receptor Tyrosine Kinase AXL in Cancer Progression.Cancers (Basel). 2016 Nov 9;8(11).

29. Morita K, Sasaki H, Furuse M, Tsukita S. Endothelial claudin: claudin-5/TMVCF constitutes tight junction strands in endothelial cells. J Cell Biol. 1999 Oct 4;147(1):185–94.

30. Ishizaki T, Chiba H, Kojima T, Fujibe M, Soma T, Miyajima H, et al. Cyclic AMP induces phosphorylation of claudin-5 immunoprecipitates and expression of claudin-5 gene in blood-brain-barrier endothelial cells via protein kinase A-dependent and -independent pathways. Exp Cell Res. 2003 Nov;290(2):275–88.

31. Fujibe M, Chiba H, Kojima T, Soma T, Wada T, Yamashita T, et al. Thr203 of claudin-1, a putative phosphorylation site for MAP kinase, is required to promote the barrier function of tight junctions. Exp Cell Res. 2004 Apr 15;295(1):36–47.

32. Yamazawa S, Shinozaki-Ushiku A, Abe H, Tagashira A, Aburatani H, Fukayama M. Gastric Cancer With Primitive Enterocyte Phenotype: An Aggressive Subgroup of Intestinal-type Adenocarcinoma. Am J Surg Pathol. 2017 Jul;41(7):989–97.

33. Reinhard K, Rengstl B, Oehm P, Michel K, Billmeier A, Hayduk N, et al. An RNA vaccine drives expansion and efficacy of claudin-CAR-T cells against solid tumors. Science. 2020 Jan 2.

34. Campbell MJ, Tonlaar NY, Garwood ER, Huo D, Moore DH, Khramtsov AI, et al. Proliferating macrophages associated with high grade, hormone receptor negative breast cancer and poor clinical outcome. Breast Cancer Res Treat. 2011 Aug;128(3):703–11.

35. Murphy LC, Seekallu SV, Watson PH. Clinical significance of estrogen receptor phosphorylation. Endocr Relat Cancer. 2011 Feb;18(1):R1–14.

36. Kato A, Hojo Y, Higo S, Komatsuzaki Y, Murakami G, Yoshino H, et al. Female hippocampal estrogens have a significant correlation with cyclic fluctuation of hippocampal spines. Front Neural Circuits. 2013;7:149.

37. Anbalagan M, Rowan BG. Estrogen receptor alpha phosphorylation and its functional impact in human breast cancer. Mol Cell Endocrinol. 2015 Dec 15;418 Pt 3:264–72.

38. Tabassum DP, Polyak K. Tumorigenesis: it takes a village. Nat Rev Cancer. 2015 Aug;15(8):473–83.

39. Cheung LWT, Hennessy BT, Li J, Yu S, Myers AP, Djordjevic B, et al. High frequency of PIK3R1 and PIK3R2 mutations in endometrial cancer elucidates a novel mechanism for regulation of PTEN protein stability. Cancer Discov. 2011 Jul;1(2):170–85.

40. Urick ME, Rudd ML, Godwin AK, Sgroi D, Merino M, Bell DW. PIK3R1 (p85a) is somatically mutated at high frequency in primary endometrial cancer. Cancer Res. 2011 Jun 15;71(12):4061–7.

41. Kandoth C, McLellan MD, Vandin F, Ye K, Niu B, Lu C, et al. Mutational landscape and significance across 12 major cancer types. Nature. Nature Publishing Group; 2013 Oct 17;502(7471):333–9.

42. Urick ME, Bell DW. Clinical actionability of molecular targets in endometrial cancer. Nat Rev Cancer. 2019 Sep;19(9):510–21.

43. Kuramoto H, Tamura S, Notake Y. Establishment of a cell line of human endometrial adenocarcinoma in vitro. Am J Obstet Gynecol. 1972 Dec 15;114(8):1012–9.

44. Cermak T, Doyle EL, Christian M, Wang L, Zhang Y, Schmidt C, et al. Efficient design and assembly of custom TALEN and other TAL effector-based constructs for DNA targeting. Nucleic Acids Res. 2011 Jul 6;39(12):e82–2.

45. Sakuma T, Ochiai H, Kaneko T, Mashimo T, Tokumasu D, Sakane Y, et al. Repeating pattern of non-RVD variations in DNA-binding modules enhances TALEN activity. Sci Rep. 2013 Nov 29;3:3379.

46. Sugimoto K, Hui SP, Sheng DZ, Nakayama M, Kikuchi K. Zebrafish FOXP3 is required for the maintenance of immune tolerance. Dev Comp Immunol. 2017 Aug;73:156–62.

47. Chiba H, Sakai N, Murata M, Osanai M, Ninomiya T, Kojima T, et al. The nuclear receptor hepatocyte nuclear factor 4alpha acts as a morphogen to induce the formation of microvilli. J Cell Biol. 2006 Dec 18;175(6):971–80.

